# Evolutionary Convergence of the Arcuate Fasciculus in Marmosets and Humans

**DOI:** 10.1101/2025.04.21.649746

**Authors:** Yufan Wang, Luqi Cheng, Deying Li, Yuheng Lu, William D. Hopkins, Chet C. Sherwood, Ting Xu, Cirong Liu, George Paxinos, Tianzi Jiang, Congying Chu, Lingzhong Fan

## Abstract

The marmoset is a highly vocal platyrrhine monkey that shares key anatomical and functional features with humans, offering insights into the evolution of brain connectivity. Although similarities in vocalization features with humans have been reported, it remains unclear whether marmosets possess an arcuate fasciculus (af) homolog. This study delineated white matter tracts in marmosets, establishing homologies with those observed in other primates, including macaques, chimpanzees, and humans. The presence of an af homolog in marmosets was confirmed by tracer and ultra-high-resolution diffusion magnetic resonance imaging datasets. We compared cortical connectivity patterns across these species and found the af in marmosets terminates in the ventral frontal cortex, with greater similarity to humans than macaques. Furthermore, we linked af connectivity with vocalization-related brain activation in both marmosets and humans. Collectively, our findings suggest that a dorsal pathway, which emerged early in marmoset evolution, has evolved convergently with humans, despite their distant phylogenetic kinship.

## Introduction

The common marmoset (*Callithrix jacchus*) is a highly vocal platyrrhine monkey that shares certain features of vocalization with humans, such as pitch perception, vocalization control, and antiphonal calling ^1–3^. The species is increasingly recognized as valuable in neuroscience and biomedical research ^4,5^, suitable for comparative analyses to elucidate the origins of evolutionary adaptations in brain structure and function along the phylogenetic tree ^6,7^. The marmoset serves as a promising comparative reference for understanding human brain evolution because of its small, less furrowed brain that facilitates neuroimaging and electrophysiological recording ^8^, along with its well-known sophisticated and distinctive social life characterized by cooperative family groups, strong pair bonds, shared parental care, and vocal communication to maintain group cohesion ^9,10^.

Recent studies have reported the capacity for increased voluntary control of vocalizations in nonhuman primates ^11,12^, suggesting a potential neuroanatomical precursor of language shared with humans ^13^. This is supported by the presence of a cortical dorsal-ventral dual stream architecture in several primate species, including marmosets. As a main component of the dorsal stream, the arcuate fasciculus (af) has been confirmed in our closest relatives, such as great apes and macaque monkeys, through non-invasive mapping of brain connections using diffusion MRI (dMRI) ^14,15^. Current views emphasize the implication of the arcuate fasciculus in human speech and language processing ^16^, with its evolutionary adaptations underpinning the complex cognitive functions observed in humans compared to nonhuman primates ^14,17–19^. However, the arcuate fasciculus homolog in marmosets, potentially linking the posterior temporal and ventral frontal regions dorsally, and the comparative connectivity patterns of this structure across species require further clarification.

In this study, we identified an arcuate fasciculus homolog in marmosets, consistent with evidence in other primates ^20–22^. The presence of an af homolog in marmosets was confirmed using tracer and ultra-high-resolution dMRI scanning data. To comprehensively explore cortical connection patterns across species, we reconstructed major white matter tracts in marmosets in a similar manner and discovered that the arcuate fasciculus connectivity in the marmoset ventral frontal cortex resembles human patterns more closely than those of macaques. The connectivity pattern of the af across species was associated with species-specific vocal communication activation patterns. We quantitatively examined these by transformed them into a common space through functional alignment, revealing a human-unique extension into the temporal cortex and a remarkable similarity between humans and marmosets in the anterior connection of the af. Our findings highlight species-specific cortical connectivity and suggest convergent evolution of the arcuate fasciculus connections in marmosets and humans, despite an early divergence in the phylogenetic tree. The logic of our analysis is outlined in Figure 1, providing an overview of the study’s methodology.

**Figure 1.**
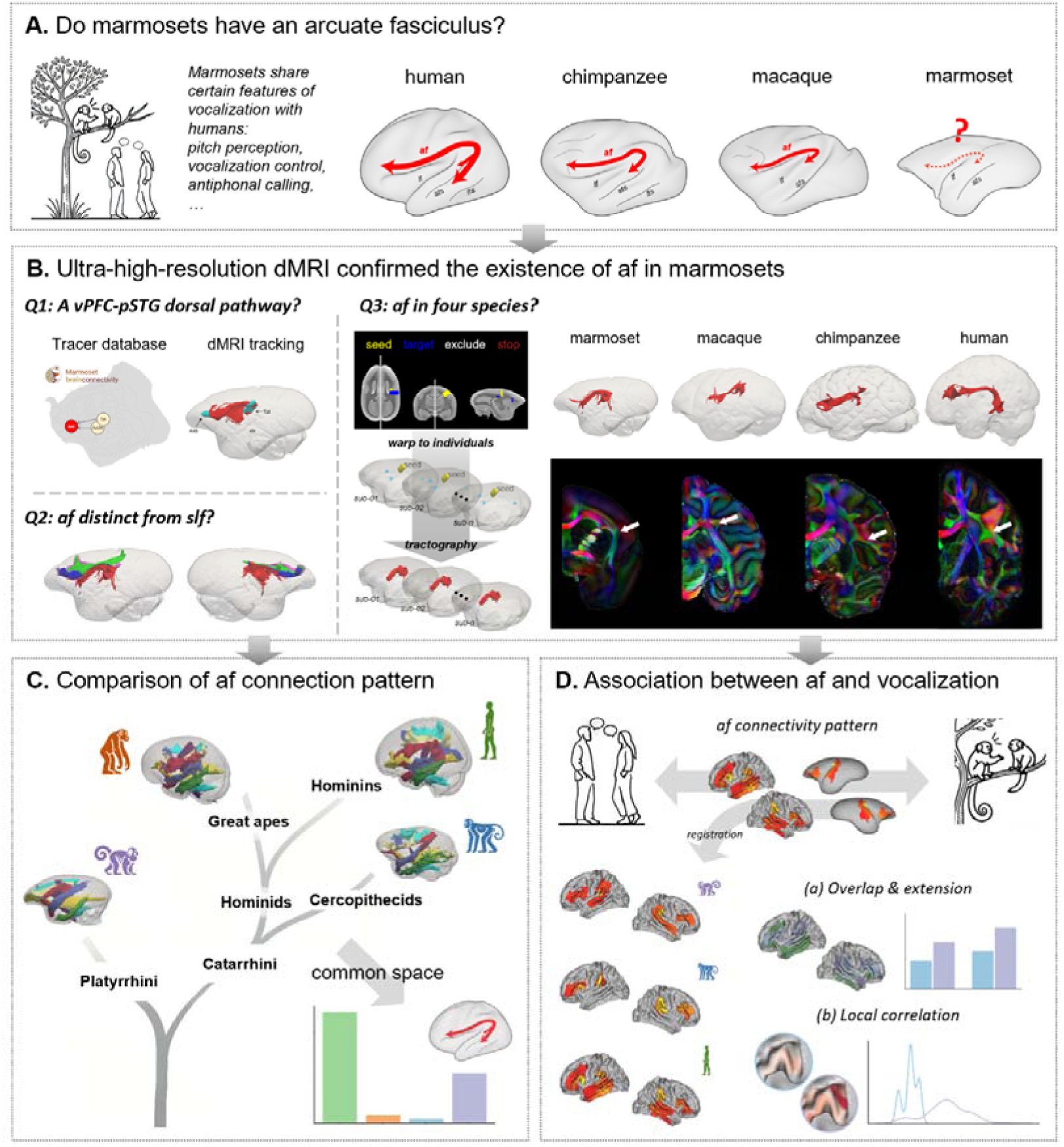
Analysis pipeline. **(A)** Given the shared vocalization features between marmosets and humans, we ask: Do marmosets have an arcuate fasciculus? **(B)** Tracer and ultra-high-resolution dMRI data were used to reveal the existence of the af in marmosets and to disambiguate it from the slf. The arcuate fasciculus homolog in marmosets was reconstructed in a manner that corresponds with those available for other primates. **(C)** Connectivity blueprints were constructed for marmosets, macaques, chimpanzees, and humans. This common space approach was further used to examine regions with significant evolutionary changes. **(D)** The connectivity pattern of the af across species was associated with vocal perception and production, and further transformed into a common space for quantitative analysis of their uniqueness.

## Results

### Marmosets have an arcuate fasciculus

Controversial is the existence in the marmoset of the homologue of the arcuate fasciculus (af), a dorsal pathway interconnecting the posterior temporal cortex and ventral frontal cortex (Figure 2A). We collected both retrograde and anterograde tracer data previously published for the marmoset brain, specifically in the Brain/MINDS Marmoset Connectivity Resource (BMCR) ^23,24^ and Marmoset Brain Connectivity Atlas (MBCA) ^25,26^. The retrograde tracing results showed connections between area 45 (A45) and the belt auditory cortex, specifically the auditory cortex caudomedial area (AuCM), and temporoparietal transitional area (Tpt) (Figure 2B). Furthermore, the anterograde neural tracer injected into A45 revealed a clear dorsal pathway above the insula (Figure 2C, left and middle), connecting with AuCM and Tpt (Figure 2C, right).

**Figure 2.**
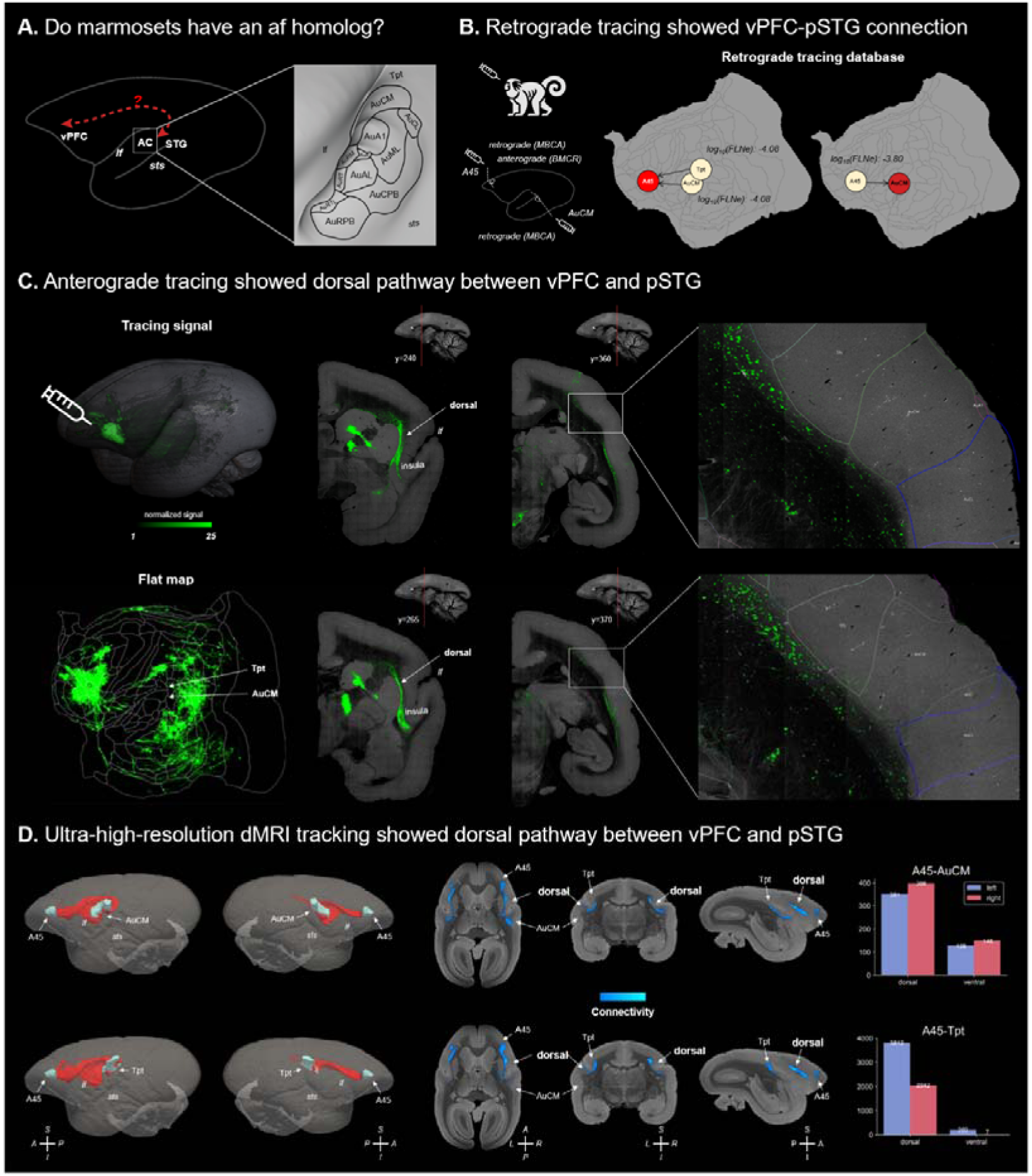
Both tracer and dMRI tracking showed a dorsal pathway between vPFC and pSTG. **(A)** Do marmosets have an arcuate fasciculus that dorsally interconnects the vPFC and pSTG? **(B)** Both retrograde and anterograde tracer data were collected from the marmoset brain. The retrograde tracing experiments showed connections between A45 and AuCM and Tpt. **(C)** The anterograde tracer injected into the A45 revealed a clear dorsal pathway above the insula, connecting the AuCM and Tpt. **(D)** A dorsal pathway connecting A45 and Tpt/AuCM was clearly observed in high-resolution dMRI images of marmosets, with this pathway being stronger than the ventral counterpart. A45, area 45; AuCM, auditory cortex caudomedial area; Tpt, temporoparietal transitional area; AC, auditory cortex; af, arcuate fasciculus; lf, lateral fissure; sts, superior temporal sulcus.

Inspired by the results from tracer experiments, we also investigated this dorsal connection using an ultra-high-resolution dMRI dataset ^27^. We set A45 in the ventral frontal cortex and the AuCM as the seed and target for diffusion tractography, and vice versa. The same procedure was conducted for A45 and the Tpt. Averaged tracking probability maps showed a dominant dorsal pathway in both hemispheres (Figure 2D), providing evidence for the existence of an af homolog in marmosets.

To reconstruct the arcuate fasciculus in marmosets, we created tractography protocols which were defined similarly to those previously established for other primates ^20–22,28^, while adjusting for the specific neuroanatomical structures of marmosets (Figure S1A). Additionally, given that the arcuate fasciculus and superior longitudinal fasciculus (slf) are often entangled, especially in post-mortem blunt dissections, we also considered the three branches of the superior longitudinal fasciculus (Figure S1B-D). In the ultra-high-resolution individual diffusion space, we reconstructed four white matter tracts (Figure 3) and identified clearly distinct pathways for each tract (Figure 3B), suggesting the disambiguation of the af from the three different branches of the slf in marmosets.

**Figure 3.**
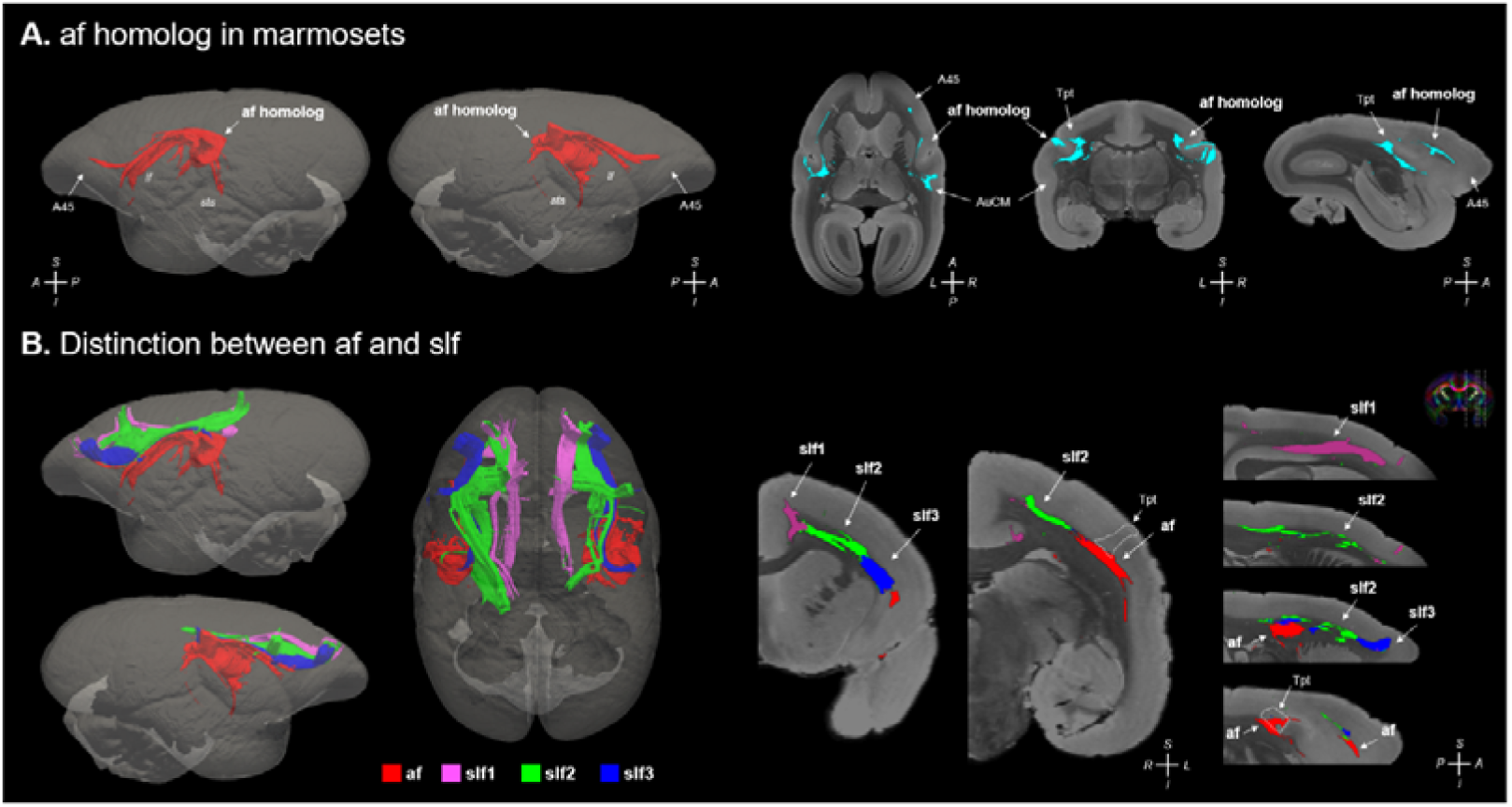
Validation of arcuate fasciculus in marmosets using an ultra-high-resolution dMRI dataset. **(A)** An arcuate fasciculus homolog was identified in marmosets. **(B)** The arcuate fasciculus was distinct from the three branches of the superior longitudinal fasciculus, as shown in high-resolution dMRI images of marmosets. A45, area 45; AuCM, auditory cortex caudomedial area; Tpt, temporoparietal transitional area; af, arcuate fasciculus; slf, superior longitudinal fasciculus; lf, lateral fissure; sts, superior temporal sulcus.

We also used high-resolution dMRI datasets from humans, chimpanzees, and macaques ^29–31^ and reconstructed the arcuate fasciculus using previously established tractography protocols. All four species showed generally consistent arched shapes that interconnect the posterior temporal and ventral frontal areas, with the most obvious difference being that the human af extends to the middle and inferior temporal lobes (Figure 4). Given the differing brain sizes across species, we resampled the diffusion data according to their brain sizes and scanning resolution, and reconstructed the af similarly. The results showed that the af was also found in the resampled data (Figure S2). These findings collectively support the existence of an arcuate fasciculus homolog in marmosets.

**Figure 4.**
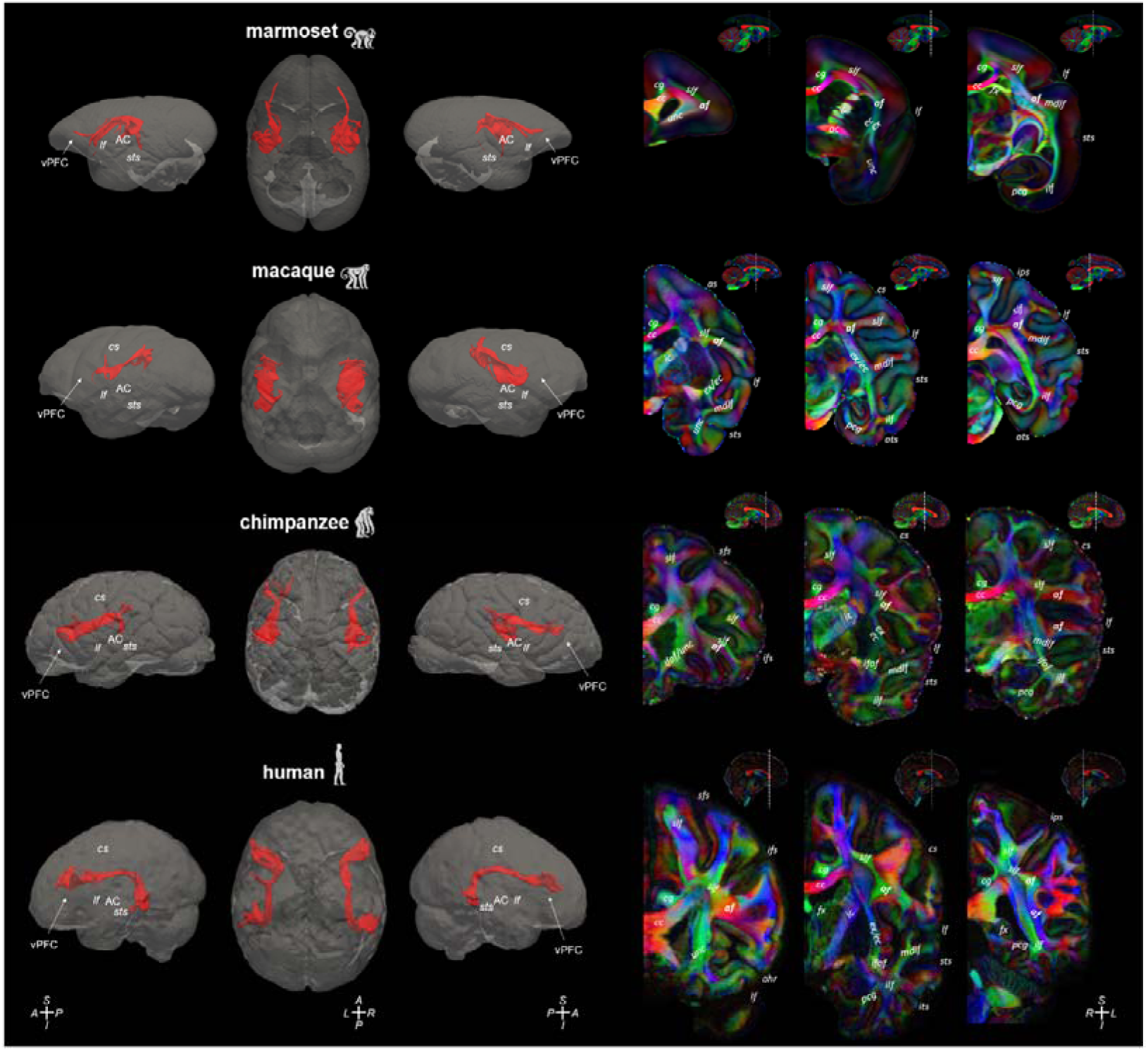
Validation of arcuate fasciculus in four species using ultra-high-resolution dMRI datasets. The arcuate fasciculus of four species was reconstructed using comparable tractography protocols. All four species showed consistent arched shapes that interconnect the posterior temporal and ventral frontal areas, with the most obvious difference being that the human af extends into the middle and inferior temporal lobes. cs, central sulcus; lf, lateral fissure; sts, superior temporal sulcus; as, arcuate sulcus; ots, occipitotemporal sulcus; ips, intraparietal sulcus; ahr, anterior ramus of lateral fissure; vPFC, ventral prefrontal cortex; AC, auditory cortex; af, arcuate fasciculus; slf, superior longitudinal fasciculus; unc, uncinate fasciculus; cg, cingulum bundle; cc, corpus callosum; ic, internal capsule; ac, anterior commissure; ec, external capsule; ex, extreme capsule; fx, fornix; mdlf, middle longitudinal fasciculus; ilf, inferior longitudinal fasciculus; pcg, parahippocampal cingulum; ifof, inferior fronto-occipital fasciculus.

### Convergent connectivity in marmoset and human arcuate fasciculus

To provide a comprehensive depiction of the cortical connectivity changes across the four species, we developed protocols to reconstruct other major white matter tracts in marmosets in a manner consistent with those available for other primates (Figure 5A, Figure S1). The tracts were described using the terminology previously employed for comparable species in the supplementary results. The robustness and generalizability of the tractography protocols were tested using an independent marmoset dataset (Brain/MINDS marmoset dataset) with varying data quality and acquisition parameters (Figure S3) ^32^.

**Figure 5.**
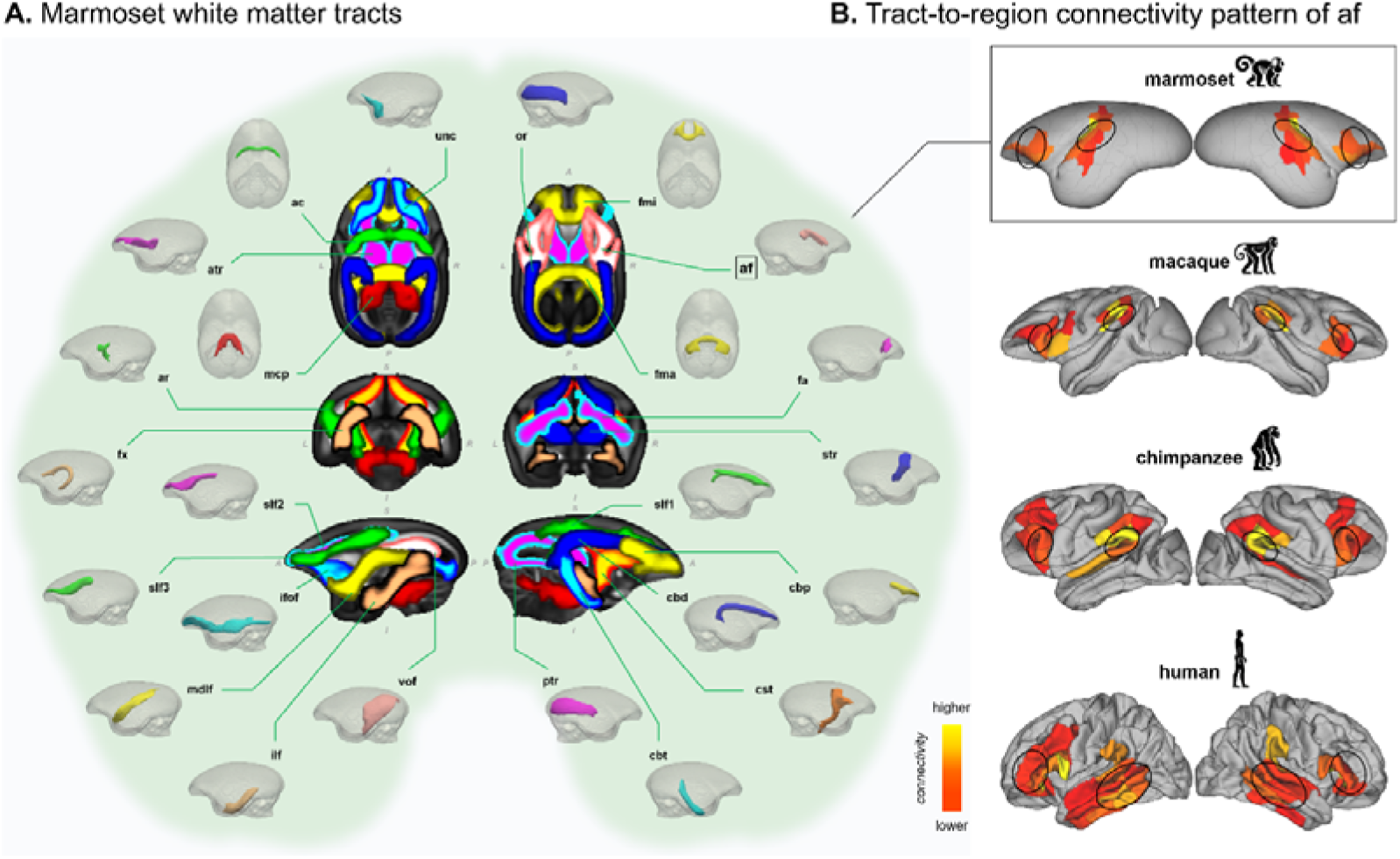
Marmoset white matter tracts and tract-to-region connectivity pattern of af. **(A)** Maximal intensity projections in horizontal, coronal, and sagittal planes of the population percentage tract atlases are shown in the center (display range = 30% - 100% of population coverage), with the 3D visualization displayed in the periphery. **(B)** The connectivity pattern of the arcuate fasciculus across marmosets, macaques, chimpanzees, and humans. af, arcuate fasciculus; slf1, superior longitudinal fasciculus I; slf2, superior longitudinal fasciculus II; slf3, superior longitudinal fasciculus III; mdlf, middle longitudinal fasciculus; ilf, inferior longitudinal fasciculus; ifof, inferior fronto-occipital fasciculus; unc, uncinate fasciculus; fa, frontal aslant tract; vof, vertical occipital fasciculus; cbd, cingulum bundle: dorsal; cbp, cingulum bundle: peri-genual; cbt, cingulum bundle: temporal; fx, fornix; ac, anterior commissure; fma, forceps major; fmi, forceps minor; mcp, middle cerebellar peduncle; cst, corticospinal tract; ar, acoustic radiation; or, optic radiation; atr, anterior thalamic radiation; str, superior thalamic radiation; ptr, posterior thalamic radiation.

To investigate the cortical connectivity patterns of the arcuate fasciculus, we counted the number of times the af tractogram reached the gray/white matter interface of each cortical region, obtaining a projection map of af connectivity for each species. All four species showed connections with the ventral frontal and premotor regions, inferior parietal regions, and superior temporal regions, but the af extended into the middle temporal regions only in humans (Figure 5B). Specifically, the af in marmosets connected to A45, proisocortical motor region (ProM), ventral area 6 (A6V), parietal area PF/PFG (PF/PFGb), Tpt, auditory belt area medial (BeltM), and other regions (Figure 5B, top).

Next, we calculated the “connectivity blueprint” by multiplying the unwrapped white matter tract matrix by the whole-brain connectivity matrix ^21^, where the rows represented each subregion’s connectivity distribution pattern. We used the connectivity blueprints for humans, chimpanzees, macaques, and marmosets to compare the connectional changes in the arcuate fasciculus across species. A modified dissimilarity measure used to quantify the difference between two probability distributions, the symmetric Kullback-Leibler (KL) divergence ^21^, was calculated to measure the dissimilarity of the regional connectivity blueprints of humans and each of the three nonhuman primates, comparing each subregion in the corresponding Brainnetome atlas. The minimum divergence for each subregion of the Human Brainnetome Atlas (HumanBNA) ^33^ was used to represent the connectivity divergence for that region, resulting in a connectivity divergence map (Figure 6A). Higher values on the divergence map indicate regions in humans with connectivity pattern that differ more from corresponding regions in the other species, i.e., it might not be represented in this close phylogenetic relative ^34^.

**Figure 6.**
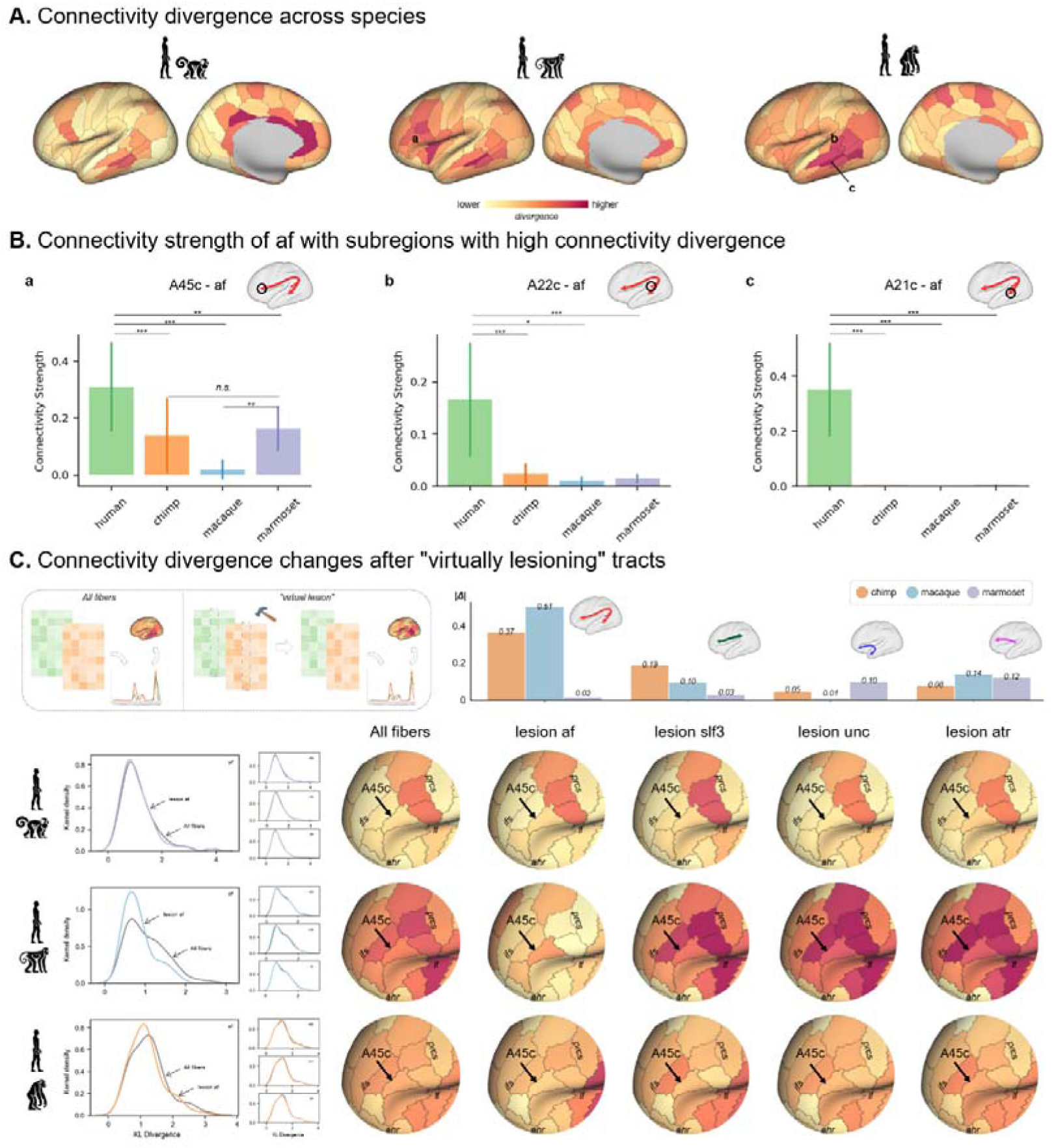
Connectivity divergence across marmosets, macaques, chimpanzees, and humans. **(A)** From left to right: connectivity divergence maps between humans and marmosets, macaques, and chimpanzees, respectively. Humans exhibit unique connectivity profiles in the middle temporal regions, lateral frontal cortex, posterior parietal cortex, and anterior cingulate cortex compared to the other three nonhuman primates. **(B)** Connectivity strength of the af with subregions showing high connectivity divergence. For example, the connection between A45c and af was weak in macaques, but stronger in marmosets and strongest in humans. **(C)** Connectivity changes after “virtually lesioning” the white matter tracts connected with A45c, including af, slf3, unc, and atr. Connectivity divergences between species changed differently across tracts, with the af causing more variable changes across cortical regions compared to the other three tracts. Focusing on the A45c, the KL divergence between humans and marmosets showed the smallest change, while macaques and chimpanzees exhibited larger changes. A45c, caudal area 45; A22c, caudal area 22; A21c, caudal area 21; af, arcuate fasciculus; slf3, superior longitudinal fasciculus III; unc, uncinate fasciculus; atr, anterior thalamic radiation; ifs, inferior frontal sulcus; prcs, precentral sulcus; lf, lateral fissure; ahr, anterior ramus of lateral fissure. *n.s.*, non-significant, * *p* < 0.05, ** *p* < 0.01, *** *p* < 0.001.

As shown in the left panel of Figure 6A and Figure S4A, the regions with the most different connectivity patterns between humans and marmosets were primarily found in the anterior and middle cingulate cortex and the dorsolateral prefrontal cortex (PFC). The regions with the most different connectivity patterns between humans and macaques were in the dorsal and medial prefrontal cortex and posterior inferior parietal lobule (IPL) (Figure 6A, middle; Figure S4B). In contrast, the regions with the most different connectivity patterns between humans and chimpanzees were located in the middle and posterior temporal lobe, posterior IPL, anterior precuneus (Pcun), insula, and inferior frontal gyrus (IFG) (Figure 6A, right; Figure S4C).

Meanwhile, minimum divergence indicates which subregion in the nonhuman primate brain shows the most similar connectivity pattern to that of humans, i.e., a “homologous” subregion defined by connectivity. This method was validated using cytoarchitectonic data to ensure cross-species correspondence ^35,36^ (Figure S5). For example, when examining human caudal area 45 (A45c), the KL divergence between its connectivity blueprint and that of each marmoset subregion was calculated. The minimum divergence was observed in area 45 in marmosets (Figure S5A), which shared similar laminar features with human area 45, including larger pyramidal cells in layer III and a well-defined granular layer IV (Figure S5B) ^37,38^, indicating cytoarchitectonic homology between the two species in this region.

We focused on brain regions connected to the arcuate fasciculus. Interestingly, we found that the connection between the af and ventral PFC (A45c) was stronger in marmosets than in macaques (*t*_human-chimp_ = 5.17, *p*_corrected_ < 0.001; *t*_human-macaque_ = 5.08, *p*_corrected_ < 0.001; *t*_human-marmoset_ = 4.19, *p*_corrected_ < 0.01; *t*_marmoset-chimp_ = 0.76, *p* = 0.45; *t*_marmoset-macaque_ = 4.85, *p*_corrected_ < 0.01; Figure 6B, a). This suggests that the connectivity of marmosets along the dorsal pathway is more similar to that in humans than in macaques, consistent with previous findings ^1^. The af extended into the temporal lobe (caudal area 22 [A22c]) and even caudally to the caudal area 21 (A21c) (a caudal area in the middle temporal regions), resulting in human-unique connectivity of the af compared to the other three nonhuman primates (A22c: *t*_human-chimp_ = 7.68, *p*_corrected_ < 0.001; *t*_human-macaque_ = 3.90, *p*_corrected_ < 0.05; *t*_human-marmoset_ = 6.62, *p*_corrected_ < 0.001; A21c: *t*_human-chimp_ = 13.52, *p*_corrected_ < 0.001; *t*_human-macaque_ = 5.70, *p*_corrected_ < 0.001; *t*_human-marmoset_ = 9.81, *p*_corrected_ < 0.001; Figure 6B, b & c). This conclusion was supported by additional macaque and marmoset data (Figure S6).

As mentioned in previous findings, the dorsal connection with the ventral PFC was more similar between humans and marmosets, a platyrrhine monkey that split from their last shared common ancestor much earlier than macaques and chimpanzees. This may suggest the marmoset’s af has continued to evolve along the phylogenetic lineage, becoming more similar to humans through convergent evolution under socio-environmental conditions that favor similarities in function. We further explored how connectivity would change after “virtually lesioning” the af in the four species (Figure 6C, top left). The connectivity divergences between species changed differently (Kolmogorov-Smirnov test: marmoset: *p* = 0.969; macaque: *p* = 0.007; chimpanzee: *p* = 0.216; Figure 6C, bottom left). Focusing on A45c, the KL divergence between humans and marmosets showed the smallest change (Δ_marmoset_=-0.02), while macaques and chimpanzees showed higher changes (Δ_macaque_ =-0.51 and Δ_chimp_ =-0.37; Figure 6C). These differences were not obvious when knocking out other white matter tracts, including the third branch of superior longitudinal fasciculus (slf3) (Δ_marmoset_ =-0.03, Δ_macaque_ =-0.10, Δ_chimp_ =-0.19), uncinate fasciculus (unc) (Δ_marmoset_ =-0.10, Δ_macaque_ =-0.01, Δ_chimp_ =-0.05), and anterior thalamic radiation (atr) (Δ_marmoset_=-0.13, Δ_macaque_=-0.14, Δ_chimp_=-0.08; Figure 6C, right). This suggests the specificity and involvement of af projections in the ventral PFC compared to other connected tracts. Meanwhile, these findings provide additional evidence for the higher similarity of the af connection with the ventral PFC between humans and marmosets.

Finally, given the shared features of vocalization between humans and marmosets, we sought to assess the involvement of the af in vocal communication in humans and marmosets. The human af is well known for its role in speech and language processing, supported by the overlap between part of its cortical connection and the functional activation of “speech” obtained from the NeuroSynth database ^39^ (Figure 7A; see also the “speech perception” and “speech production” in Figure S7). Specifically, auditory cortex and more posterior superior temporal cortex were activated during speech perception, while the inferior frontal gyrus, ventral premotor, and motor cortex were activated during speech production.

**Figure 7.**
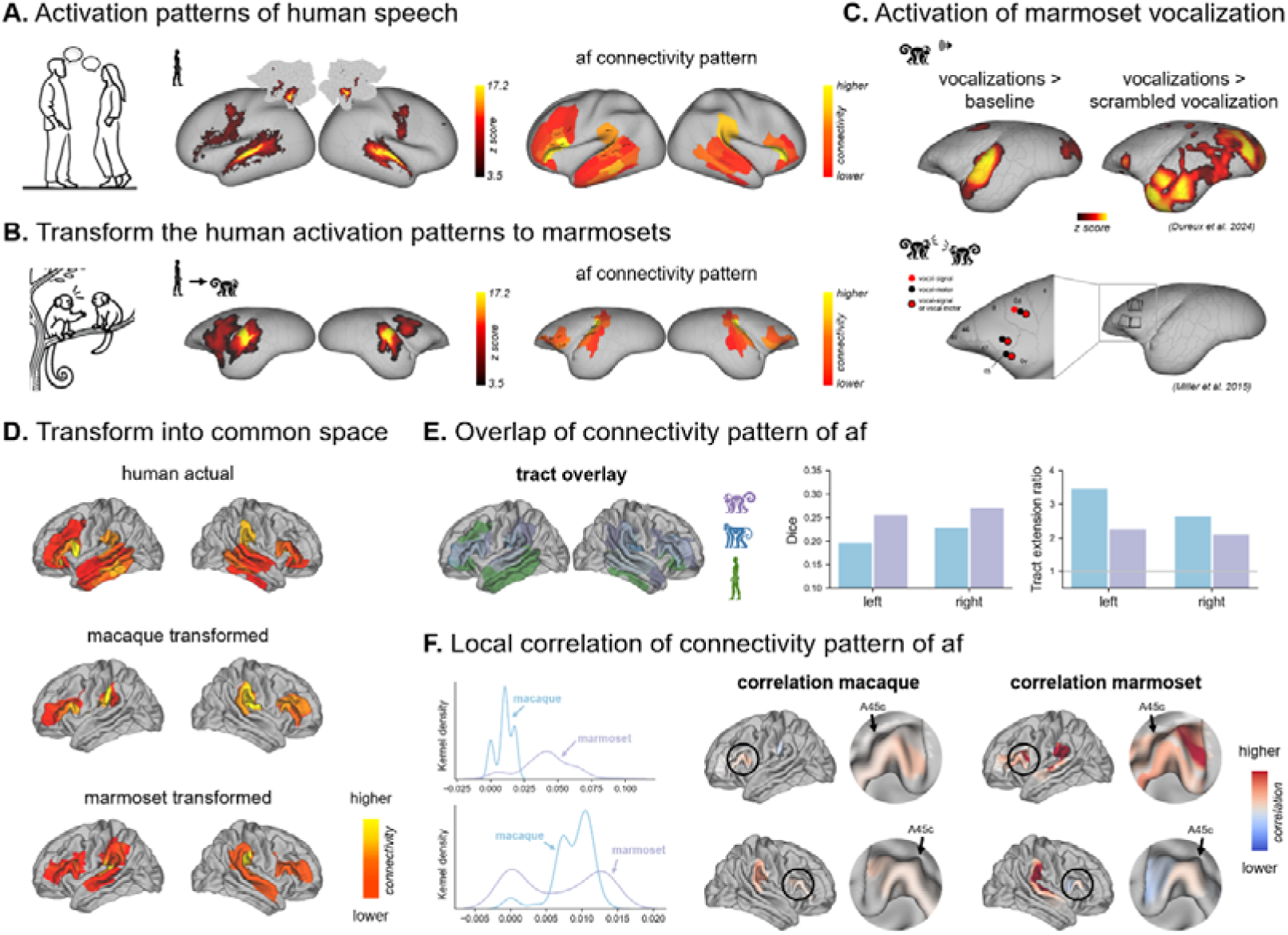
Comparison of af connectivity patterns across species. **(A)** Activation patterns of human speech and their association with the connectivity patterns of the af. **(B)** The transformed activation patterns of human speech in the marmoset cortex. **(C)** Activation patterns associated with marmoset vocal communications. **(D)** The af connectivity patterns transformed from macaques and marmosets using functional alignment techniques. **(E)** Binarized af connectivity patterns across the three species, along with quantitative Dice coefficients and tract extension ratios for each map. **(F)** Weighted correlation maps of the actual human map and the transformed macaque and marmoset maps. The left panel shows the distribution of weighted correlation values in the A45c.

We transformed these activation maps to marmoset cortex based on functional-alignment techniques ^40,41^ (Figure 7B), which showed overlap with the connectivity pattern of the af in marmosets, including A6V, and others. Although whole-brain imaging of marmosets during vocal communication is difficult to obtain, previous fMRI and electrophysiological experiments provide important evidence of the involvement of these cortical regions in vocalization perception ^42^ and natural vocal communication ^43^ (Figure 7C).

We further quantitatively evaluated the similarity of connectivity patterns of the af across species by transforming them into a common space using the same functional-alignment techniques (Figure 7D). Note the chimpanzees were not considered here because their functional data were unavailable. We first calculated the Dice coefficient to assess the overlap similarity between the actual and transformed tract connection map, and a “tract extension ratio” to assess how much of the human tract projections extend into areas where other species do not ^18^. Marmosets showed the highest tract connection overlap with humans compared to macaques (left: *Dice*_macaque_ = 0.196, *Dice*_marmoset_ = 0.255; right: *Dice*_macaque_ = 0.227, *Dice*_marmoset_ = 0.271; Figure 7E). The tract extension ratios for the three species showed a similar degree, with all values greater than 1, indicating that the af in humans extended further into more brain areas during evolution (left: *Ratio*_macaque_ = 3.457, *Ratio*_marmoset_ = 2.251; right: *Ratio*_macaque_ = 2.627, *Ratio*_marmoset_ = 2.089; Figure 7E).

To further localize where the af showed similar connections across species, we calculated the weighted local correlation map between the actual and transformed tract connection map, where higher values indicated brain areas with both the tract of humans and the other species showing a termination. Focusing on the ventral PFC (A45c), the results showed that marmosets exhibited higher similarity with humans than macaques (median correlation in the left hemisphere: *r*_macaque_ = 0.0107, *r*_marmoset_ = 0.0426; right: *r*_macaque_ = 0.0097, *r*_marmoset_ = 0.0119; Figure 7F).

## Discussion

In this study, we identified an arcuate fasciculus homolog in marmoset brains that corresponds to those in other primates. We also found that marmosets exhibited a more similar connection pattern of the arcuate fasciculus with humans in the ventral frontal cortex, which was further supported by quantitative analyses after transformation into a common space. Our results suggest the presence of an earlier-emerged dorsal pathway in marmosets that convergently evolved aspects of its connectivity with humans.

Previous studies have largely focused on the Catarrhini, neglecting potential evolutionary changes in the Platyrhini after their ancestors split. By leveraging species-equivalent tractography protocols, which are informed by anatomical insights into reconstructing the arcuate fasciculus in human, chimpanzee, and macaque brains ^20^, we identified a dorsal fiber pathway that connects the posterior temporal and ventral frontal regions, traversing through the inferior parietal and ventral premotor areas, while also being distinct from the three branches of the superior longitudinal fasciculus, suggesting the presence of the long-suspected arcuate fasciculus homolog in marmosets. Additionally, tract-tracing experiments showed that area 45 in the ventral frontal cortex receives monosynaptic auditory input from the belt auditory cortex ^25,26^ and also sends information to the superior temporal cortex and posterior parietal cortex ^23,24^. These dMRI and tracer observations of the putative arcuate fasciculus might push back the emergence of this dorsal pathway further than the split from a common ancestor with marmosets, approximately 35 million years ago ^14,44^. The present data provide further evidence for the auditory dorsal pathway among primate species, a pathway previously observed in tractography following seeding in auditory regions ^1,27^, supporting species-specific vocalizations during social behaviors.

Most interestingly, unlike the weak connection to the ventral frontal cortex in macaques, marmosets exhibited a stronger connection, more similar to that in humans, suggesting that the dorsal pathway in marmosets may have evolved independently in closer alignment with humans than with macaques after the marmoset lineage diverged ^1^. The similarity in the ventral frontal cortex between humans and marmosets may help explain the similarity in the perception of vocalizations in marmosets and speech in humans ^45^. Production and use of vocalizations in primates show similar spectral characteristics and neural circuits with human speech ^46^, particularly in the prefrontal and premotor cortex of marmosets, which has been found to be associated with the production of social communication signals across the marmoset vocal repertoire ^47^. Another complex process associated with both vocal perception and production, previously unreported in nonhuman primates_—_voluntary vocal control, the modification of vocal production according to the acoustic environment—was observed in marmosets, allowing them to avoid acoustic interference and maintains effective communication ^48,49^.

Our data raise the question of whether these changes in the arcuate fasciculus are unique to marmosets or shared among other platyrrhines, as no studies have yet examined this feature in other platyrrhine species, such as capuchin monkeys, howler monkeys, or others. It is also unknown whether these species share an ancestral trait which undergoes parallel evolutionary modifications independently. This calls for broader comparative studies to map af connectivity across more primate species, even those with different vocalization patterns, and to integrate genetic and developmental perspectives for deeper insights.

Comparable vocal communication complexity may have exerted similar selective pressure on brain connectivity in humans and marmosets, both species having evolved in environments requiring adaptability and intricate social interactions, which might have driven the enhancement of auditory-motor integration. The highly cooperative social structures favored stronger sensorimotor integration for real-time social coordination or even predictive social processing ^50–52^, reinforcing af-like pathways beyond vocal communications.

In contrast to the findings in the ventral frontal cortex, the arcuate fasciculus showed relatively higher connectivity differences within the middle and posterior temporal cortex, along with observed hemispheric differences between species, underscoring a potential human-specific adaptation ^53,54^. These adaptations likely played a role in the evolution of language in the human lineage, particularly in enabling the development of more complex linguistic abilities, such as speech processing ^54,55^. Despite these differences, vocal behavior among primate species provides a window into the foundational elements of human speech, suggesting an early evolutionary origin of some of its components. This behavior was further refined in human language to enhance social communication ^13^.

In addition to its relevance to speech, this study provided a detailed marmoset white matter tract atlas comparable to those of other primate species, facilitating cross-species investigations of similarities and differences in fiber pathways. As an important primate model, the common marmoset shares key brain organizational features with larger primates, making it feasible to identify marmoset white matter tracts based on comparative evidence from other primates ^27,56^. Although previous research has presented a detailed delineation of white matter pathways using ultra-high-resolution dMRI datasets ^27^, that mapping was limited to a single *ex vivo* subject, reducing its generalizability to larger cohorts and its utility for direct quantitative comparison with other species. The literature is now complemented by the robust and reproducible tractography protocols we used, protocols grounded in prior anatomical knowledge and registered to individual space to reconstruct white matter tracts consistently across subjects while accounting for individual variability. The established population-based marmoset white matter tract atlas will serve as a resource for comparative research.

In conclusion, we employed dMRI to map marmoset white matter tracts, identifying an arcuate fasciculus homolog and its convergent similarities with humans in the ventral frontal cortex. This provided an opportunity to extend comparative research to more species and reveal general organizational principles of primate brains. Future exploration that also take mammals and birds into consideration could offer new insights into how intelligence evolved from divergent neural structures ^57^. Although this study preliminarily investigated the functional association of af connectivity, future comparative analyses combining electrophysiological recordings, behavioral, or lesion studies could provide direct evidence linking af connectivity to vocal or cognitive functions, supporting the investigation of cognitive specializations in humans and nonhuman primates.

## Methods

### MRI Data acquisition and preprocessing

To facilitate an intuitive comparison across datasets, we summarize the scanning parameters of the diffusion datasets used in this study, including spatial resolution and diffusion directions, in Table S1 (ultra-high-resolution datasets) and S2 (lower-resolution datasets).

### Marmoset dataset 1: Marmoset Brain Mapping (MBM)

Marmoset (*Callithrix jacchus*; *n* = 24, 17 males; mean age, 4.17±1.88 y; age range, 1.88-9.00 y) MRI scans were obtained from a data archive of the Marmoset Brain Mapping Project (https://marmosetbrainmapping.org/) ^58^. The experimental procedures followed policies established by the US Public Health Service Policy on Humane Care and Use of Laboratory Animals. All procedures were approved by the Animal Care and Use Committee (ACUC) of the National Institute of Neurological Disorders and Stroke, National Institutes of Health.

All the marmosets underwent a 3–4-week acclimatization protocol, as previously described ^58,59^. After completing the training, they were properly acclimated to lying in the sphinx position in an MRI-compatible cradle with their heads comfortably restrained with 3D-printed anatomically conforming helmets.

The marmosets were scanned in a 7T/30 cm horizontal MRI (Bruker, Billerica, USA) equipped with a 15 cm customized gradient set capable of 450 mT/m gradient strength (Resonance Research Inc., Billerica, USA) and an 8-channel phased-array RF coil custom-built for marmosets^60^. T2-weighted structural images were collected at a 0.25 × 0.25 × 0.5 mm resolution (TR = 6000 ms, TE = 9 ms, flip angle = 90°, FOV = 28 × 36 mm, matrix size = 112 × 144, 38 axis slices, slice thickness = 0.5 mm). Multishell diffusion MRI data were collected using a 2D diffusion-weighted spin-echo EPI sequence with the following parameters: TR = 5.1 s, TE = 38 ms, number of segments = 88, FOV = 36 × 28 mm, matrix size = 72 × 56, slice thickness = 0.5 mm, a total of 400 DWI images for two-phase encodings (blip-up and blip-down) and each had 3 *b* values (8 *b* = 0, 64 *b* = 1000 s/mm^2^, and 128 *b* = 2000 s/mm^2^). The multishell gradient sampling scheme was generated using the Q-shell sampling method.

The preprocessing of the *in vivo* diffusion MRI dataset incorporated denoising, correcting for eddy currents-and EPI-induced distortions using pairs of diffusion data acquired with opposite phase encoding (blip-up and blip-down), and merging the preprocessed pairs into one dataset. The nonlinear spatial registration from the individual space to the DTI template of MBMv3 space ^61^ was carried out using ANTs, which were then used to transform the landmarks defined on the template into individual space for tractography. Voxel-wise estimates of the fiber orientation distribution were calculated using bedpostx ^62^, allowing for three fiber orientations for the dataset. **Marmoset dataset 2: Brain/MINDS Marmoset Brain MRI Dataset (Brain/MINDS)**

Another cohort of adult marmosets was obtained from the Brain/MINDS Marmoset Brain MRI Dataset (https://dataportal.brainminds.jp/marmoset-mri-na216) ^32,63^ by retaining only the adult ones. Our study included 110 adult marmosets with both structural and *in vivo* diffusion-weighted images (50 males; mean age, 4.38±2.47 y; age range, 1.73-10.02 y). All procedures were approved by the Animal Experiment Committees of the RIKEN Center for Brain Science (CBS) and conducted following the Guidelines for Conducting Animal Experiments of RIKEN CBS.

The marmosets were scanned in a 9.4-T BioSpec 94/30 unit (Bruker Optik GmbH, Ettlingen, Germany) and a transmit and receive coil with an inner diameter of 86□mm. T1-weighted structural images were collected using a magnetization-prepared rapid gradient echo (MP-RAGE) with the following parameters: TR = 6000 ms, TE = 2 ms, flip angle = 12°, number of averages = 1, inversion time = 1600 ms, voxel size = 0.27 × 0.27 × 0.54 mm. T2-weighted structural images were collected using rapid acquisition with relaxation enhancement (RARE), with the following parameters: TR = 4000 ms, TE = 22 ms, RARE factor = 4, flip angle = 90°, number of averages = 1, voxel size = 0.27 × 0.27 × 0.54 mm. Multishell diffusion MRI data were collected using a spin-echo EPI sequence with the following parameters: TR = 3 s, TE = 25.6 ms, number of segments = 6, voxel size = 0.35 × 0.35 × 0.7 mm, a total of 94 DWI images (4 *b* = 0, 30 *b* = 1000 s/mm^2^, and 60 *b* = 3000 s/mm^2^). The preprocessing of the dataset was the same with the MBM dataset.

### Marmoset dataset 3: High-resolution marmoset data

A formalin-fixed brain sample from one male common marmoset (4.5 y) with ultra-high-resolution MRI scanning was used ^27^. Details of procedures before MRI scanning can be found in previous publications ^27^. The brain sample was scanned on a 7T/30-cm horizontal bore MRI spectrometer (Bruker Biospin) with a 30-mm inner diameter quadrature millipede coil (ExtendMR). Multishell diffusion MRI data were collected using a 3D diffusion-weighted multi-shot spin-echo EPI sequence with the following parameters: TR = 200 ms, TE = 29 ms, number of averages = 1, number of segments = 88, FOV = 38 × 29.76 × 29.76 mm, matrix size = 474 × 372 × 372, resolution = 80 μm isotropic, a total of 204 DWI images with four *b* values (8 *b* = 0, 6 *b* =30 s/mm^2^, 64 *b* = 2400 s/mm^2^, and 126 *b* = 4800 s/mm^2^). The diffusion MRI data of *ex vivo* brain samples had been preprocessed using the DIFFPREP of TORTOISE as in the previous study ^27^.

### Human dataset 1: Human Connectome Project (HCP)

Data from 40 healthy human adults (17 males) were selected. These were the 40 subjects that were utilized in the construction of the Human Brainnetome Atlas ^33^ as well as the ones available from the S1200 subjects release of the Human Connectome Project (HCP) database (http://www.humanconnectome.org/) ^64^. All the scans and data from the individuals included in the study had passed the HCP quality control and assurance standards.

The scanning procedures and acquisition parameters were detailed in previous publications ^65^. In brief, T1w images were acquired with a 3D MPRAGE sequence on a Siemens 3T Skyra scanner equipped with a 32-channel head coil with the following parameters: TR = 2400 ms, TE = 2.14 ms, flip angle = 8°, FOV = 224×320 mm^2^, voxel size = 0.7 mm isotropic. DWI images were acquired using single-shot 2D spin-echo multiband echo planar imaging on a Siemens 3 Tesla Skyra system (TR = 5520 ms, TE = 89.5 ms, flip angle = 78°, FOV = 210×180 mm). These consisted of three shells (*b*-values = 1000, 2000, and 3000 s/mm^2^) with 90 diffusion directions isotropically distributed among each shell and six *b* = 0 acquisitions within each shell at a spatial resolution of 1.25 mm isotropic voxels.

The diffusion-weighted images were preprocessed using FSL ^66^. Following preprocessing, voxel-wise estimates of the fiber orientation distribution were calculated using bedpostx ^62^, allowing for three fiber orientations for the human dataset.

### Human dataset 2: High-resolution human data

A whole-brain *in vivo* diffusion MRI dataset acquired at 760 µm isotropic resolution was used ^30^. The data were acquired on the MGH-USC 3T Connectom scanner equipped with high-strength gradient (G_max_□=□300 mT/m), a custom-built 64-channel phased-array coil and personalized motion-robust head stabilizer, using the recently developed SNR-efficient simultaneous multi-slab imaging technique termed gSlider-SMS. Advanced parallel imaging reconstruction with ghost-reduction algorithm (Dual-Polarity GRAPPA) and reversed phase-encoding acquisition were also performed to mitigate ghosting artifacts and image distortion. The data was totally acquired at 1260 q-space points including 420 directions at *b* = 1000 s/mm^2^ and 840 directions at *b*□=□2500□s/mm^2^ across the 9 sessions, and was employed by an optimized preprocessing pipeline.

### Chimpanzee dataset 1: National Chimpanzee Brain Resource (NCBR)

We obtained 46 adult chimpanzees (18 males) with T1w and DWI data from the National Chimpanzee Brain Resource (NCBR, http://www.chimpanzeebrain.org/). The data were acquired using previously described procedures at the Emory National Primate Research Center (ENPRC) on a 3T MRI scanner under propofol anesthesia (10 mg/kg/h) ^67^. All procedures followed protocols approved by ENPRC and the Emory University Institutional Animal Care and Use Committee (IACUC, approval no. YER-2001206). All data were obtained before the 2015 implementation of U.S. Fish and Wildlife Service and National Institutes of Health regulations governing research with chimpanzees. All chimpanzee scans were completed by the end of 2012; no new data were acquired for this study.

T1w images were collected at an 0.7 × 0.7 × 1 mm resolution. DWI images were acquired using a single-shot spin-echo echo-planar sequence for 60 diffusion directions (b = 1000 s/mm^2^, TR = 5900 ms; TE = 86 ms; 41 slices; 1.8 mm isotropic resolution). DWI images with phase-encoding directions (left-right) of opposite polarity were acquired to correct susceptibility distortion. Five *b* = 0 s/mm^2^ images were also acquired with matching imaging parameters for each repeat of a set of DWI images.

The diffusion-weighted images were preprocessed similarly using FSL ^66^. Following preprocessing, voxel-wise estimates of the fiber orientation distribution were calculated using bedpostx ^62^, allowing for two fiber orientations for the chimpanzee dataset due to the *b*-value in the diffusion data.

### Chimpanzee Dataset 2: High-resolution chimpanzee data

A whole-brain chimpanzee dMRI dataset was acquired at 500□µm isotropic voxel size on a preclinical Bruker Biospec 94/30 MRI system at 9.4T (Paravision 6.0.1), using a G_max_□=□300 mT/m gradient system and a 154-mm transmit-receive quadrature radiofrequency coil (Bruker BioSpin). ^29^. Details of procedures before MRI scanning can be found in previous publications ^29^. Diffusion MRI data were collected using a segmented 3D EPI spin-echo sequence with the following parameters: TR = 1,000 ms, TE = 58.9 ms, matrix size = 240□×□192□×□144, no partial Fourier, no parallel acceleration, EPI segmentation factor of 32 and EPI-Readout-BW of 400□kHz, 55 DWI images with *b* = 5000 s/mm^2^. Three interspersed *b*□=□0 images without diffusion-weighting and an additional *b*□=□0 volume were acquired with reversed-phase encoding direction for correction.

### Macaque dataset 1: TheVirtualBrain (tvb)

Data from eight adult rhesus macaque monkeys (all males) were obtained from TheVirtualBrain ^68^. All surgical and experimental procedures were approved by the Animal Use Subcommittee of the University of Western Ontario Council on Animal Care (AUP no. 2008–125) and followed the Canadian Council of Animal Care guidelines. Surgical preparation and anesthesia as well as imaging acquisition protocols have been previously described ^68^. Images were acquired using a 7T Siemens MAGNETOM head scanner. Two diffusion-weighted scans were acquired for each animal with each scan having an opposite phase encoding in the superior-inferior direction at 1 mm isotropic resolution, allowing for the correction of susceptibility-related distortion. The data were acquired with 2D EPI diffusion for five animals, and a multiband EPI diffusion sequence was used for the remaining three animals. In all cases, the data were acquired with *b* = 1000 s/mm^2^, 64 directions, and 24 slices. A 3D T1w image was also collected for each animal (128 slices, resolution: 0.5 mm isotropic).

The diffusion-weighted images were preprocessed similarly using FSL ^66^. Following preprocessing, voxel-wise estimates of the fiber orientation distribution were calculated using bedpostx ^62^, allowing for two fiber orientations for the macaque dataset due to the *b*-value in the diffusion data.

### Macaque dataset 2: UC-Davis macaque data (ucd)

Data from nineteen adult rhesus macaque monkeys (all females) were available from PRIME-DE (https://fcon_1000.projects.nitrc.org/indi/indiPRIME.html) ^69^. The neuroimaging experiments and associated procedures were performed at the California National Primate Research Center under protocols approved by the University of California, Davis Institutional Animal Care and Use Committee ^70^. Images were acquired on a 3T Siemens Skyra scanner with four-channel clamshell coil. Two diffusion-weighted scans were acquired for each animal with opposite phase encoding, allowing for the correction of susceptibility-related distortion. The data were acquired with *b* = 1600 s/mm^2^, 60 directions. T1w images were collected for each animal with resolution at 0.3 mm isotropically. The preprocessing of the dataset was the same with the macaque tvb dataset.

### Macaque dataset 3: High-resolution macaque data

An ultra-high-resolution postmortem macaque dMRI dataset was acquired at 200 μm isotropic voxel size on a 7 Tesla small animal MRI system equipped with 650 mT/m Resonance Research gradient coils ^31^. Details of procedures before MRI scanning can be found in previous publications^31^. Diffusion MRI data were collected using a 3D spin-echo pulse sequence with the following parameters: TR = 100 ms, TE = 32.1 ms, matrix size = 384□×□288□×□256, 30 DWI images with *b* = 4000 s/mm^2^ and a *b* = 0 volume were acquired.

### Neuronal tract-tracing data in marmosets

#### Dataset 1: Marmoset Brain Connectivity Atlas (MBCA)

We included retrograde neuronal tracer data from the publicly available Marmoset Brain Connectivity Atlas (MBCA, https://www.marmosetbrain.org/) ^25,26^. All experiments conformed to the Australian Code of Practice for the Care and Use of Animals for Scientific Purposes and were approved by the Monash University Animal Experimentation Ethics Committee.

The dataset comprises 143 retrograde tract-tracing experiments conducted in 52 marmosets (31 males; mean age, 2.5 y; age range, 1.4-4.6 y), injected with six different fluorescent retrograde tracers. The strength of the tract-tracing from the labeled neurons (source) to the injection area (target) was quantified by the Fraction of Labeled neurons (extrinsic; FLNe).

#### Dataset 2: Brain/MINDS Marmoset Connectivity Resource (BMCR)

We also included anterograde neuronal tracer data from the Brain/MINDS Marmoset Connectivity Resource (BMCR, https://dataportal.brainminds.jp/marmoset-connectivity-atlas) ^23,24^. All experimental procedures were carried out following the National Institute of Health Guide for the Care and Use of Laboratory Animals and the Japanese Physiological Society’s Guiding Principles for the Care and Use of Animals in the Field of Physiological Science and were approved by the Experimental Animal Committee of RIKEN.

This dataset includes anterograde tracer image data from 52 marmosets (19 males; mean age, 5.9 y; age range, 2.3-10.8 y). For 19 of these animals, the anterograde tracer data are complemented by retrograde tracer data. Injections were targeted to 21 brain regions in the left hemisphere of the marmoset prefrontal cortex.

#### Connections between regions of interest

To investigate whether a dorsal connection exists between the ventral prefrontal cortex and the posterior temporal cortex, we selected cortical regions based on the Paxinos atlas of the marmoset brain ^35^, including area 45 (A45) anteriorly, and auditory cortex caudomedial area (AuCM), and temporoparietal transitional area (Tpt) posteriorly. We assessed the connection strength between each pair of regions of interest, i.e., A45 and AuCM, and A45 and Tpt, using retrograde data from the MBCA.

Notably, the anterograde tracer image data were mapped into a 3D reference image space ^24^, enabling visual examination of the pathways of neuronal connections. We selected data from Monkey #R01_0110, in which the injection site was located in A45.

### Tractography protocols

#### Tractography protocols of white matter tracts

The protocols for reconstructing white matter tracts were defined similarly to those previously used for other primates ^20–22,28^, albeit adjusted for the neuroanatomical structures of marmosets. All protocols included a midsagittal section as an exclusion mask to isolate the tracts within one hemisphere, except for the anterior commissure, forceps major, forceps minor, and middle cerebellar peduncle, which link brain regions across hemispheres. Here, we provide descriptions of the arcuate fasciculus and superior longitudinal fasciculus, while those for other major white matter tracts can be found in the supplementary methods.

The marmoset arcuate fasciculus was reconstructed with a seed underneath the PF/PFG, a posterior target axially posterior to the terminus of the lateral fissure, and an anterior target at the level of the ventral premotor cortex, posterior to the ventral prefrontal cortex.

For the three branches of the superior longitudinal fasciculus, a coronal mask in the white matter underneath the primary motor cortex and primary somatosensory cortex was used as the seed, along with two target masks. Anteriorly, target masks for the first, second, and third branches of the slf were coronal sections through the BA8/BA9, BA9/BA46, and BA45/BA47, respectively. Posteriorly, target masks for the first, second, and third branches of the slf were coronal sections through the PE, IP/PG, and PF/PFG, respectively. For each subcomponent, the seed was positioned in alignment with the posterior target and moved anteriorly into the primary motor/somatosensory regions. The exclusion mask consisted of an axial mask underneath the parietal cortex, an axial mask excluding the subcortical cortex, and a coronal mask preventing ventral longitudinal tracts.

#### Tractography to reconstruct white matter tracts

Probabilistic diffusion tractography was performed using FSL’s probtrackx2 ^62^. The seed, target, exclusion, and stop masks, defined in volumetric standard space, were warped to individual space for tractography. The tractography was run in two directions (seed to target and target to seed) with the following parameters: each seed voxel was sampled 10,000 times, with a curvature threshold of 0.2, and a number of steps of 3200. The step length was set to one-fourth of voxel size according to each dataset. The resulting tractograms were normalized by dividing by the waypoint number and then warped back to template space. The normalized tractograms were thresholded at 0.005 for individuals and averaged to create a group-level tractogram for each tract. The tracts were reconstructed in MNI152 standard space for humans, Yerkes29 space for chimpanzees, F99 space for macaques, and MBMv3 space for marmosets.

#### Tractography between regions of interest

To be consistent with the tracer results, we selected cortical regions based on the Paxinos atlas of the marmoset brain ^35^. For each pair of regions of interest, i.e., A45 and AuCM, and A45 and Tpt, we conducted diffusion tractography using one region as the seed and the other as the target, and vice versa, to generate two sets of tracking probability maps. The two probability maps were averaged into a single map to represent the final connection. Dorsal and ventral waypoints were defined separately to compare the relative strength of the pathways based on the tracking results.

#### Construction of connectivity patterns of tracts

We counted the number of times the tractogram of white matter tract reached the gray/white matter interface of each cortical region, obtaining a projection map of the tract for each species. These projection maps were normalized by the size of seed mask (in voxels) for each subject of each species to allow valid cross-species comparisons.

### Connectional comparison across four species

#### Construction of connectivity blueprints

We performed probabilistic tractography using FSL’s probtrackx2 ^62^ to map the whole-brain connectivity. Specifically, the white surface was set as the seed region, tracking to the rest of the brain with the ventricles removed. The pial surface served as a stop mask to prevent streamlines from crossing sulci. Each vertex was sampled 10,000 times (10,000 trackings) based on the orientation probability model for each voxel, with a curvature threshold of 0.2 and a number of steps of 3200. The step length was set to one-fourth of voxel size according to each dataset, resulting in a (*whole-surface vertices*) × (*whole-brain voxels*) matrix for further analysis.

The homologous white matter tracts of four species were reconstructed using XTRACT following protocols ^20,22^. The tractography data were then vectorized into a (*tracts*) × (*whole-brain voxels*) matrix and multiplied by the whole-brain connectivity matrix, deriving a (*tracts*) × (*whole-surface vertices*) matrix, i.e., the connectivity blueprint ^21^. The matrix rows showed each vertex’s connectivity distribution pattern.

We further averaged the connectivity profiles of the vertices within subregions of atlas of each species. Specifically, we averaged the connectivity profiles of the vertices within each subregion of the Human Brainnetome Atlas (HumanBNA) ^33^, Chimpanzee Brainnetome Atlas (ChimpBNA) ^34^, Macaque Brainnetome Atlas (MacBNA) ^71^, and Marmoset Brain Mapping Atlas (MBM) ^72^ to form the regional connectivity blueprints across subjects for humans, chimpanzees, macaques, and marmosets, respectively. Note that the middle cerebellar peduncle (mcp) does not project to the cortex, and was excluded from constructing the blueprints. So, the final number of tracts = 44.

#### Connectivity divergence between species

We used connectivity blueprints for each species to explore the connectivity divergence across species ^21,34^. We calculated the symmetric Kullback Leibler (KL) divergence for each pair of subregions in the MBM and HumanBNA and searched for the minimal value for each subregion in the HumanBNA. Specifically, for an example subregion in the HumanBNA, the connectivity blueprint represents how it is connected to each white matter tract. Using the KL divergence measure, this connectivity profile was compared with those of all marmoset subregions. The subregion with the connectivity blueprint most similar to the human one, i.e., with the smallest KL divergence, was selected, and this KL divergence value was assigned to the human subregion. A higher value on this cross-species divergence map means that the subregion has a more dissimilar connectivity profile that is absent in marmosets. The same procedure was conducted between humans and chimpanzees and humans and macaques.

#### Comparison of connectivity strength of af

We compared the individual connectivity strength of af with subregions across four species. Specifically, we focused on caudal area 45 (A45c), caudal area 22 (A22c), and caudal area 21 (A21c). Connectivity probability was compared between species using a two-sample t test, with significance determined after correcting for multiple comparisons using Bonferroni correction. **“Virtually lesioning” the tracts in connectivity blueprints**

To examine the effect of specific white matter tract on the connectivity divergence, we proposed a “virtually lesioning” approach by setting the connectivity probability values of the tract to zero and recalculating the minimum KL divergence. The difference in connectivity divergences between species was examined using the Kolmogorov-Smirnov test.

### Functional alignment and tract connection comparison

#### Functional alignment across species

The cross-species functional alignment among marmosets, macaques, and humans was used ^40,41^. Specifically, homologous landmarks in the three species were identified for extracting matched functional components using the joint-embedding approach, which were then input into multimodal surface matching (MSM) ^73^. The macaque-human transformation was used as an intermediate step to construct the cortical transformation between marmosets and humans. Chimpanzees were not considered here because their functional data was unavailable.

#### Alignment of functional activation

To investigate the association between functional activation by speech and the connectivity patterns of the af, we selected terms such as “speech,” “speech perception,” and “speech production” from the NeuroSynth database ^39^ (https://neurosynth.org/) and projected them onto the surface. The activation maps were transformed into the marmoset surface for comparison with the connectivity patterns of af in marmosets.

#### Comparison of connectivity patterns of af in a common space

We compared the connectivity patterns of af across species in a common human space. We transformed the connectivity patterns of marmosets and macaques to human space. We calculated Dice coefficients and tract extension ratio to quantify the amount of overlap in the tract maps ^18^. The Dice coefficient was computed for the binarized actual human tract map and the map transformed from the other species. The tract extension ratio was calculated as the ratio of the number of vertices covered by the human tract map to the number of vertices covered by both the human and the other tract map.

To intuitively and quantitatively compare tract connections between humans and nonhuman primates, we calculated weighted local correlation maps of the human map and those transformed from other species. Specifically, the local correlation map was computed using a sliding window around every vertex on the sphere, which was adjusted to up-weight the brain areas where the tract is represented on the surface. Higher values in the weighted correlation map indicated brain areas where both the human tract and the tract from the other species terminate.

## Data availability

The tractography protocols in standard space for producing marmoset white matter tracts and their surface projection data are available online (https://github.com/FANLabCASIA/MarmosetWM). The tractography protocols for humans, chimpanzees, and macaques are available at https://git.fmrib.ox.ac.uk/rmars/chimpanzee-tractography-protocols.

The MBM dataset and the high-resolution single marmoset subject data are available at https://marmosetbrainmapping.org/data.html. The Brain/MINDS dataset is available at https://dataportal.brainminds.jp/marmoset-mri-na216. The human data are available from the Human Connectome Project (https://db.humanconnectome.org/). The high-resolution single human subject data is available at https://doi.org/10.5061/dryad.nzs7h44q2. The chimpanzee data are available at the National Chimpanzee Brain Resource (http://www.chimpanzeebrain.org/). The high-resolution single chimpanzee subject data is available from the Evolution of Brain Connectivity Project (https://doi.org/10.17617/3.O5XSI9). The TVB macaque data are available at https://doi.org/10.18112/openneuro.ds001875.v1.0.3. The UC-Davis macaque data are available at https://fcon_1000.projects.nitrc.org/indi/PRIME/ucdavis.html. The high-resolution single macaque subject data is from a previously shared resource. The Brain/MINDS Marmoset Connectivity Resource is available at https://dataportal.brainminds.jp/marmoset-tracer-injection. The Marmoset Brain Connectivity Atlas is available at https://analytics.marmosetbrain.org/graph.html. The human and marmoset Nissl-stained sections are from a previously shared resource.

## Code availability

The HCP-Pipeline can be found at https://github.com/Washington-University/HCPpipelines, and the NHP-HCP-Pipeline can be found at https://github.com/Washington-University/NHPPipelines. The neuroimaging preprocessing tools used are freely available, including FreeSurfer v6.0 (http://surfer.nmr.mgh.harvard.edu/) and FSL v6.0.7 (https://fsl.fmrib.ox.ac.uk/fsl/fslwiki). The brain maps were presented using MRIcroGL v20 (https://www.nitrc.org/projects/mricrogl) and Workbench v1.5.0 (https://www.humanconnectome.org/software/connectome-workbench). The tracts were visualized using ITK-SNAP 4.0.1 (http://www.itksnap.org/) and Paraview 5.11.0

(https://www.paraview.org/). The code for analysis used in this study is available at https://github.com/FANLabCASIA/MarmosetWM.

## Acknowledgments

We thank Kadharbatcha Saleem for valuable discussion. This work was partially supported by STI2030-Major Projects (Grant No. 2021ZD0200203), the Natural Science Foundation of China (Grant Nos. 82072099, 82202253, 62250058), the China Postdoctoral Science Foundation (2022M722915, 2024M761725), and Guangxi Science and Technology Base and Talent Special Project (Grant No. AD22035125). Data were provided in part by the National Chimpanzee Brain Resource (supported by NIH NS092988, NIH HG011641, NIH AG067419, NIH AG087945, NSF EF-2021785, and NSF DRL-2219759), the Duke Center for In Vivo Microscopy NIH/NIBIB (P41 EB015897), Human Connectome Project from WU-Minn Consortium (Principal Investigators: David Van Essen and Kamil Ugurbil; 1U54MH091657) funded by the 16 NIH Institutes and Centers that support the NIH Blueprint for Neuroscience Research and by the McDonnell Center for Systems Neuroscience at Washington University. The authors thank Tianlei Zhang for his help in visualizing the results, and appreciate the English language and editing assistance of Rhoda E. and Edmund F. Perozzi, PhDs.

