## supplementary information for "Evolutionary Convergence of the Arcuate Fasciculus in Marmosets and Humans"

This supplemental information contains

Supplementary methods

Supplementary results

Supplementary figures S1-S7

Supplementary tables S1-S2

References

### Supplementary methods

#### Tractography protocols

The protocols were defined as similarly as possible to those previously defined for other primates ^1-4^ albeit adjusted for the neuroanatomical structures of marmosets. All protocols included a midsagittal section as an exclusion mask for isolating the tracts within one of the two hemispheres, except for the anterior commissure, forceps major, forceps minor, and middle cerebellar peduncle, which link brain regions between the left and right hemispheres.

**Association tracts**

**Arcuate fasciculus (af)** The marmoset arcuate fasciculus was reconstructed with a seed underneath the PF/PFG, a posterior target axially posterior to the terminus of the lateral fissure, and an anterior target at the level of the ventral premotor cortex, posterior to the ventral prefrontal cortex.

**Superior longitudinal fasciculus I/II/III (slf 1/2/3)** For the three branches of the superior longitudinal fasciculus, a coronal mask in the white matter underneath the primary motor cortex and primary somatosensory cortex was used as the seed along with the two target masks. Anteriorly, target masks for the first, second, and third branches of the slf were coronal sections through the BA8/BA9, BA9/BA46, and BA45/BA47, respectively. Posteriorly, target masks for the first, second, and third branches of the slf were coronal sections through the PE, IP/PG, and PF/PFG, respectively. For each subcomponent, the seed was positioned in alignment with the posterior target while moved anteriorly into the primary motor/somatosensory regions. The exclusion mask consisted of an axial mask underneath the parietal cortex, an axial mask excluding the subcortical cortex, and a coronal mask preventing ventral longitudinal tracts.

**Middle longitudinal fasciculus (mdlf)** The seed and target of the mdlf were coronally placed in the anterior and posterior parts of the dorsal temporal lobe, respectively. The target was located just anterior to the posterior terminus of the lateral fissure. The exclusion mask consisted of an axial mask through the brainstem, a coronal mask through the fornix, an axial mask through the cingulum, and a coronal mask through the frontal cortex. The seed and target mask of the ilf served as additional exclusion masks.

**Inferior longitudinal fasciculus (ilf)** The seed and target of the ilf were coronally placed in the anterior and posterior parts of the ventral temporal lobe, respectively. The exclusion mask consisted of an axial mask through the brainstem, a coronal mask through the fornix, an axial mask through the cingulum, a coronal mask through the frontal cortex, and a coronal mask through the PF/PFG in the parietal cortex. In addition, the seed mask of the mdlf served as an exclusion mask.

**Inferior fronto-occipital fasciculus (ifof)** The seed of the ifof was a coronal plane through the anterior part of the occipital cortex, and the target was a coronal plane through the frontal cortex anterior to the genu of the corpus callosum. An exclusion mask was used to restrict all fibers running through the extreme/external capsule.

**Uncinate fasciculus (unc)** The seed for the unc was placed in the white matter where the temporal and frontal cortex separate. The target was placed in the ventral extreme/external capsule. An exclusion mask between the seed and target was used to force the curve. An additional mask was placed coronally to prevent fibers from leaking longitudinally to the temporal lobe.

**Frontal aslant tract (fa)** The fa seed was placed sagittally in the white matter of the ventral prefrontal cortex, the target was placed axially in the white matter of the dorsal prefrontal cortex. A posterior coronal exclusion mask prevented leakage into longitudinal fibers.

**Vertical occipital fasciculus (vof)** The axial seed and target masks were placed in the lateral part of the occipital white matter dorsally and ventrally, respectively. A coronal exclusion mask was set just posterior to the corpus callosum to prevent leakage into anterior-posterior tracts.

**Limbic tracts**

**Cingulum bundle (cbd, cbp, cbt)** The cingulum bundle was segmented into three sections based on the targets that were connected to the fibers. The dorsal part (cbd) was seeded just above the posterior part of the corpus callosum and was targeted at the start of the genu of the corpus callosum. A sagittal mask in the anterior limb of the internal capsule was used as an exclusion to prevent leakage into the temporal lobe. The peri-genual part (cbp) was seeded in the dorsal genu of the corpus callosum and targeted at the ventral sub-genual callosum. A coronal plane at the level of the rostral end of the lateral fissure was used as an exclusion to prevent leakage into the cbd. The temporal part (cbt) was seeded in the posterior part of the temporal lobe and targeted posterior to the amygdala. An exclusion mask was used to prevent leaking into the fornix. Two stop masks were placed posteriorly and anteriorly to the seed and target masks, respectively.

**Fornix (fx)** The fornix was seeded in the body of the fornix and targeted in the hippocampus. The exclusion mask consisted of a coronal plane anterior to the occipital cortex to prevent leakage into posterior tracts and a sagittal mask around the midline at the level of the anterior tip of the thalamus to prevent lateral propagation to the anterior limb of the internal capsule. A stop mask was added axially in the ventral temporal lobe to prevent leakage into the cingulum.

**Commissure tracts**

**Anterior commissure (ac)** The seed was placed at the midline of the anterior commissure, and the target was placed in the white matter between the globus pallidus and the putamen. An exclusion mask was placed superior to the seed, and stop masks were placed posterior to the seed and target masks to prevent leakage via the rest of the basal ganglia and were placed axially at the level of the extreme/external capsule to prevent leakage via the ventral pathway.

**Forceps major and forceps minor (fma, fmi)** The seed and target mask for the forceps major were defined as coronal sections through the anterior part of the occipital lobe. The exclusion mask consisted of a coronal plane through the pons to prevent longitudinal fibers and a sagittal mask to confine the area to the occipital cortex. The seed and target mask for the forceps minor were defined as coronal sections through the frontal lobe anterior to the corpus callosum. The exclusion mask consisted of a coronal plane located at the anterior third of the corpus callosum level and of a midsagittal mask to prevent posterior projections.

**Middle cerebellar peduncle (mcp)** The middle cerebellar peduncle was seeded in the cerebellar white matter with a target in the opposite hemisphere. An exclusion mask was placed sagittally between the cerebellar hemispheres and axially through the thalamus.

**Projection tracts**

**Corticospinal tract (cst)** The corticospinal tract was seeded in the pons with a target covering the premotor, motor, and somatosensory cortices. The exclusion mask consisted of two coronal planes to exclude tracking into the prefrontal and occipital cortex and two axial planes to restrict tracking to the cerebral peduncle of the midbrain and cerebellar peduncles.

**Acoustic and optic radiations (ar, or)** The acoustic and optic radiations were seeded from the medial geniculate nucleus (MGN) and lateral geniculate nucleus (LGN) of the thalamus, respectively. A target was placed in the auditory core area for acoustic radiation, and the exclusion mask consisted of two coronal planes, anterior and posterior to the thalamus. The target mask for optic radiation covered a coronal section through the anterior part of the calcarine fissure. Exclusion masks were placed axially through the brainstem, coronally posterior to the LGN to select fibers curling around dorsally, and anterior to the seed to prevent leaking into longitudinal fibers.

**Anterior, superior, and posterior thalamic radiations (atr, str, ptr)** The anterior, superior, and posterior thalamic radiations connect the thalamus to the frontal lobe, primary motor/somatosensory cortex, and occipital lobe, respectively. The anterior thalamic radiation was seeded in the anterior part of the thalamus with a target in the anterior thalamic peduncle. An exclusion mask was placed posterior to the thalamus, and a coronal plane was used to prevent leakage via the cingulum. The superior thalamic radiation was seeded using the superior half of the thalamus, targeted in the primary motor cortex and primary somatosensory cortex. The exclusion mask consisted of two coronal planes to exclude tracking into the prefrontal and occipital cortex. An axial plane ventral to the thalamus was used as a stop mask. The posterior thalamic radiation was seeded in the posterior part of the thalamus with a coronal target in the occipital lobe. Exclusion masks were placed anterior and inferior to the thalamus.

#### Robustness across datasets

To explore the robustness of the protocols across data of varying quality and acquisition parameters, we used another dataset of 110 adult marmosets, i.e., the Brain/MINDS dataset ^5,6^, to reconstruct the white matter tracts. We then compared tract-atlases and intersubject variability between the two datasets. Specifically, a group of marmosets from the Brain/MINDS dataset, matched for sex with the MBM dataset, was selected for comparison. Tract atlases were compared by correlating each tract between the two datasets. The tractography results assessed intersubject variability as the average tract-wise correlations between subject pairs within and across cohorts. The similarity was assessed using the Pearson correlation coefficient between the subjects' normalized tract density maps in MBMv3 template space with a threshold of 0.5%. The correlation coefficients were averaged across the tracts for each subject pair. Significance between within- and across-cohort correlations was obtained via Mann-Whitney U test.

### Supplementary results

#### White matter tract atlas of the marmoset brain

**Association tracts**

Conventionally, the dorsal longitudinal tracts connecting the frontal lobe with the parietal and posterior temporal cortices are formed by the arcuate fasciculus (af) and three branches of the superior longitudinal fasciculus (slf). Although the nomenclature of these tracts has differed between various species and in spite of inconsistent recognition of the arcuate fasciculus in marmosets, we were able to reconstruct these tracts using a protocol that is comparable with that used for humans, chimpanzees, and rhesus macaques ^1^ by which all four tracts have been identified using dMRI tractography in these larger brained species.

In our findings, the marmoset arcuate fasciculus extended from the ventral prefrontal cortex to the posterior parietal cortex and curved into the posterior superior temporal gyrus, reaching anteriorly to the auditory areas. The superior longitudinal fasciculus I (slf1) extended from the dorsal parietal area to the dorsal prefrontal cortex. The superior longitudinal fasciculus II (slf2) ran medially to the superior longitudinal fasciculus (slf3), connecting frontal areas with intraparietal and ventral parietal areas, and the slf3 reached the ventral prefrontal areas and extended posteriorly to the ventral parietal cortex.

The marmosets' middle longitudinal fasciculus (mdlf) extended from the superior temporal rostral area through the superior temporal gyrus and reached the parietal lobe. Ventrally, the inferior longitudinal fasciculus (ilf) passed through the inferior temporal area and posteriorly extended to the inferior lateral occipital cortex. The inferior fronto-occipital fasciculus (ifof) connected the occipital cortex and prefrontal cortex rostrocaudally via the extreme/external capsule, running medial to the mdlf and ilf through the temporal lobe ^7^. The uncinate fasciculus (unc) extended from the temporopolar region and curved into the inferior and orbital frontal cortex.

The frontal aslant tract (fa) connects the ventrolateral prefrontal cortex with the dorsal frontal cortex ^8,9^. In marmosets, the fa extended from the ventral frontal cortex to area 8 in the dorsal frontal cortex superiorly. The vertical occipital fasciculus (vof) in marmosets ran dorsoventrally throughout the anterior part of occipital lobe, consistent with its location in other primates.

**Limbic tracts**

The cingulum bundle was reconstructed by combining the results from dorsal, peri-genual, and temporal sections. The cingulum bundle started from the parahippocampal area, through the medial posterior temporal lobe, coursing rostrocaudally superior to the corpus callosum, and terminating in the medial prefrontal cortex. The fornix (fx) started from the medial temporal lobe, which was connected to the mammillary bodies and reached the hypothalamus.

**Commissure tracts**

The anterior commissure (ac) crossed the midline, connecting the amygdala and temporal lobes between the two hemispheres. The middle cerebellar peduncle (mcp) originated from the pontine nuclei and travelled to the opposite cerebellar hemisphere. The forceps major (fma) and forceps minor (fmi), which are components of the corpus callosum, run through the splenium and the genu, respectively. The forceps major streamlines were mainly concentrated in the parieto-occipital cortex, while the forceps minor streamlines reached the medial prefrontal cortex and frontal pole.

**Projection tracts**

The corticospinal tract (cst) was a collection of axons descending from the primary motor/somatosensory cortex to the spinal cord. The streamlines reached the primary motor cortex and primary somatosensory cortex in marmosets, consistent with other primates.

The acoustic (ar) and optic radiations (or) arose from the medial and lateral geniculate nucleus, respectively. The ar streamlines reached the auditory area, while the or streamlines were concentrated on the occipital pole in the marmosets. The anterior (atr), superior (str), and posterior thalamic radiations (ptr) connected the thalamus to the frontal lobe, the primary motor/somatosensory cortex, and the occipital cortices, respectively.

#### Robustness of tractography protocols across datasets

We tested the robustness and generalizability of the tractography protocols using a variety of data quality and acquisition parameters on an independent marmoset dataset (Brain/MINDS marmoset dataset) ^5^. The demographics of two marmoset datasets were shown in Figure S3A and S3B.

We compared the tract atlases and the intersubject variability between the two datasets. Strong similarity was observed between the average spatial correlation across tracts in the group atlases (*μ* = 0.81, *σ* = 0.07; Figure S3C). Intersubject similarity was relatively high and consistent within the datasets, albeit with a slightly higher variance in the Brain/MINDS dataset (MBM, *μ* = 0.86, *σ* = 0.01; Brain/MINDS, *μ* = 0.75, *σ* = 0.03; *p* = 8×10^-92^ for comparison between within-MBM and within-MINDS; Figure S3D). Although the cross-cohort comparison showed a lower correlation than the within-cohort comparison (*μ* = 0.72, *σ* = 0.03; *p* = 1×10^-123^ for comparison between MBM-MINDS and within-MBM and *p* = 5×10^-45^ for comparison between MBM-MINDS and within-MINDS; Figure S3D), they were still comparable given the differences in data quality and age of the subjects in the two cohorts (MBM: mean age = 4.17±1.88 y, age range = 1.88-9.00 y; Brain/MINDS: mean age = 4.38±2.47 y, age range = 1.73-10.02 y). The above analyses demonstrated the feasibility and fidelity of tract reconstructions using a standard protocol with different data acquisition sources.

### Supplementary figures

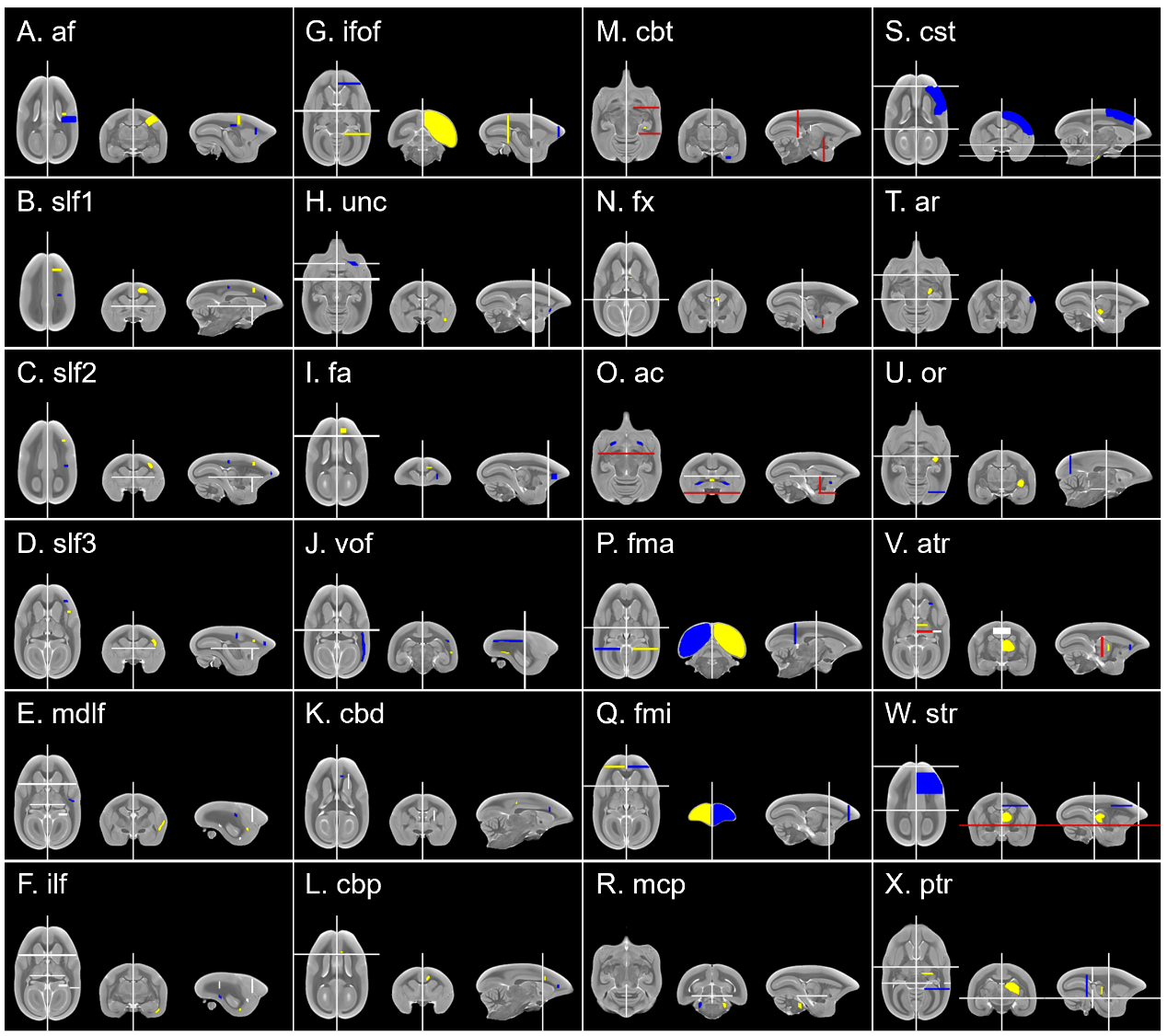

**Figure S1. Marmoset tractography protocols.** Seed (yellow), target (blue), exclusion (white) and stop (red) masks are displayed for left hemisphere in radiological convention. af, arcuate fasciculus; slf, superior longitudinal fasciculus; mdlf, middle longitudinal fasciculus; ilf, inferior longitudinal fasciculus; ifof, inferior fronto-occipital fasciculus; unc, uncinate fasciculus; fa, frontal aslant tract; vof, vertical occipital fasciculus; cbd, cingulum bundle: dorsal; cbp, cingulum bundle: peri-genual; cbt, cingulum bundle: temporal; fx, fornix; ac, anterior commissure; fma, forceps major; fmi, forceps minor; mcp, middle cerebellar peduncle; cst, corticospinal tract; ar, acoustic radiation; or, optic radiation; atr, anterior thalamic radiation; str, superior thalamic radiation; ptr, posterior thalamic radiation.

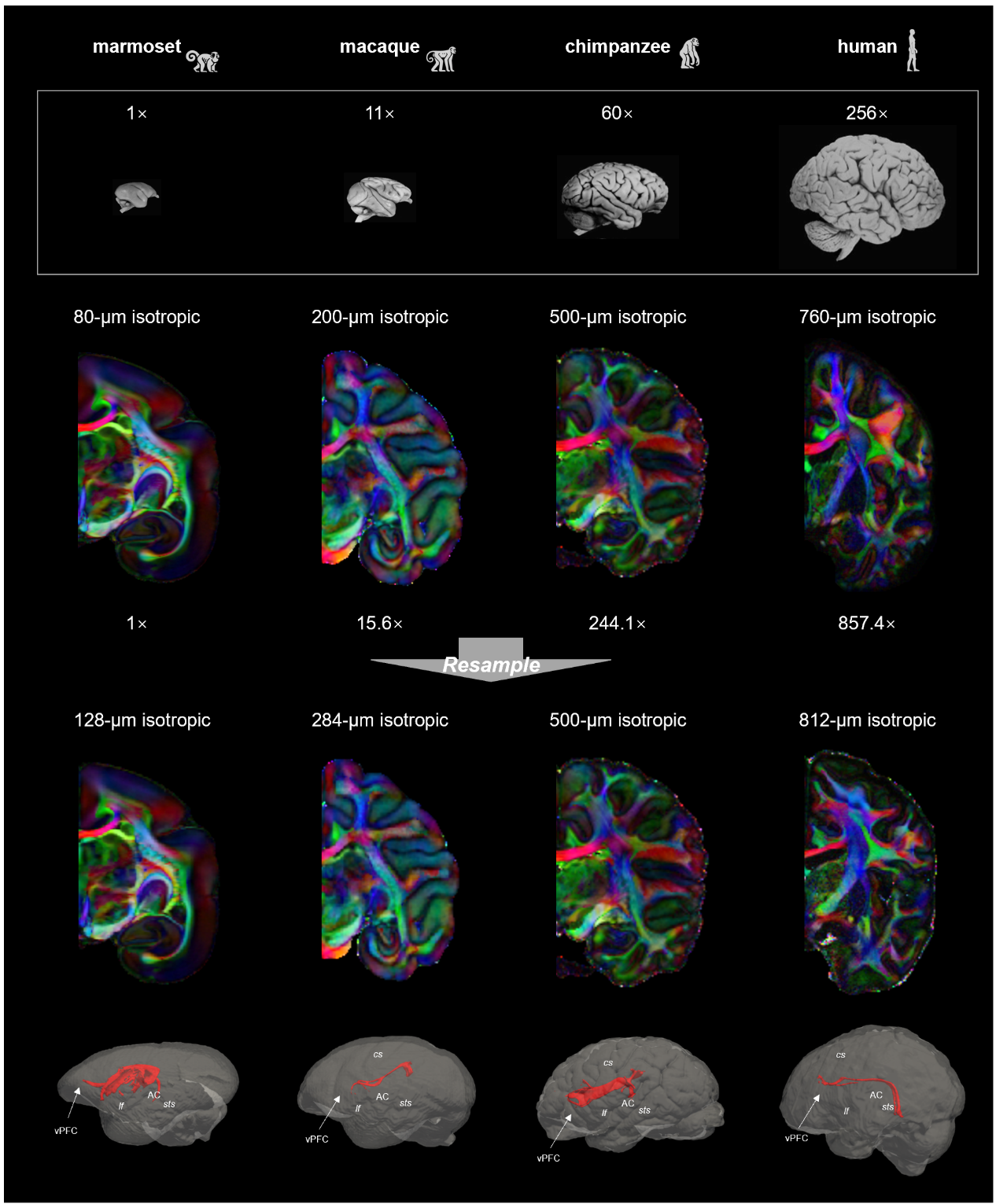

**Figure S2. Reconstruction of arcuate fasciculus in the resampled diffusion data of four species.** As the brain size differ across species, we resampled the data according to their brain size and scanning resolution. The arcuate fasciculus was reconstructed using the same way. The results showed that the af was also found in the resampled data, suggesting the stability and robustness of our tracking methods and results.

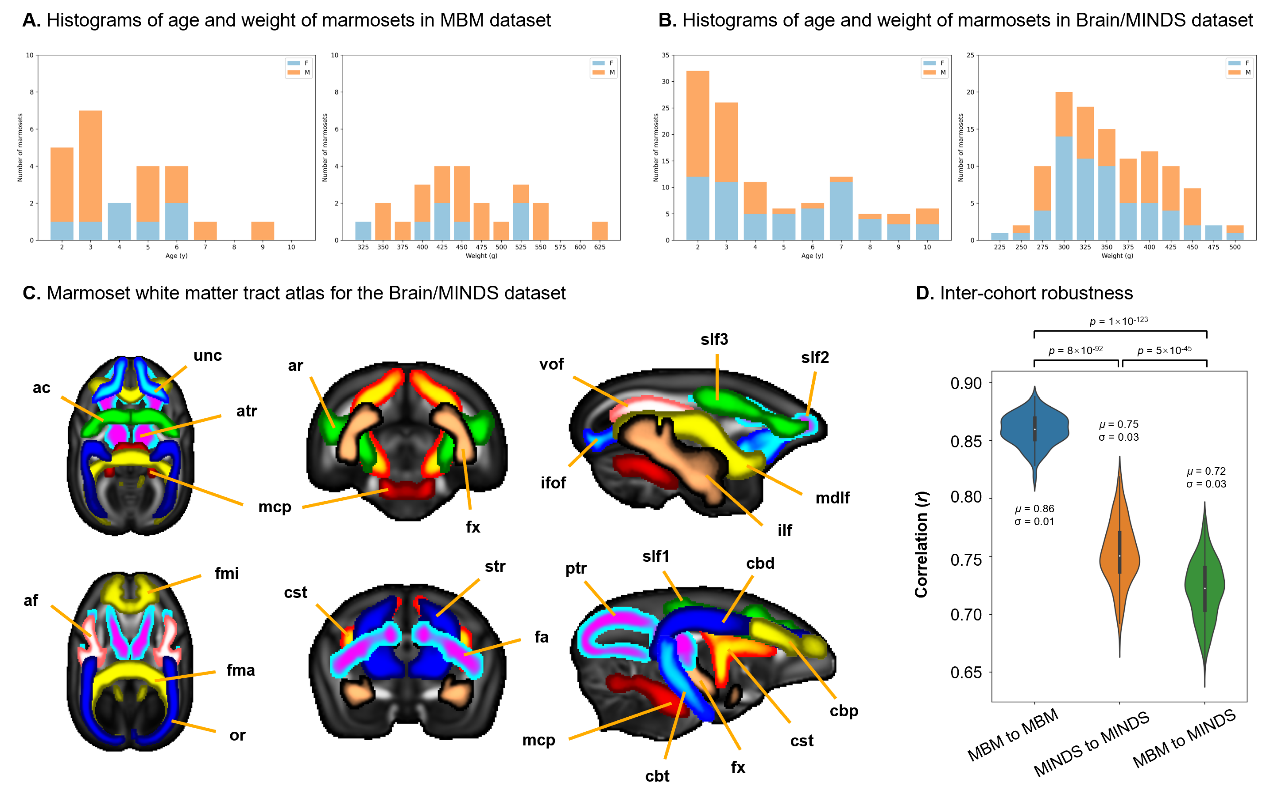

**Figure S3. Histograms of age and weight of marmosets and robustness of tractography protocols against datasets. (A)** Histograms of age and weight of marmosets according by sex in MBM dataset. **(B)** Histograms of age and weight of marmosets according by sex in Brain/MINDS dataset. **(C)** The marmoset white matter tract atlas for the Brain/MINDS dataset. Horizontal, coronal, and sagittal maximal intensity projections of the population percentage tract atlases are shown (display range = 30% - 100% of population coverage). **(D)** Summary of inter-cohort robustness. Intersubject variability was assessed as the average tract-wise correlations in the tractography results between subject pairs within and across cohorts. The intersubject variability was relatively consistent across the datasets, with greater intersubject similarity within than across groups. Correlations were calculated on normalized tract density maps with a threshold of 0.5%. *μ* was the median of the correlations across subject pairs and *σ* was the standard deviation. Significance was obtained via Mann-Whitney U test. Corrected *p*-value is 0.05/3 = 0.017.

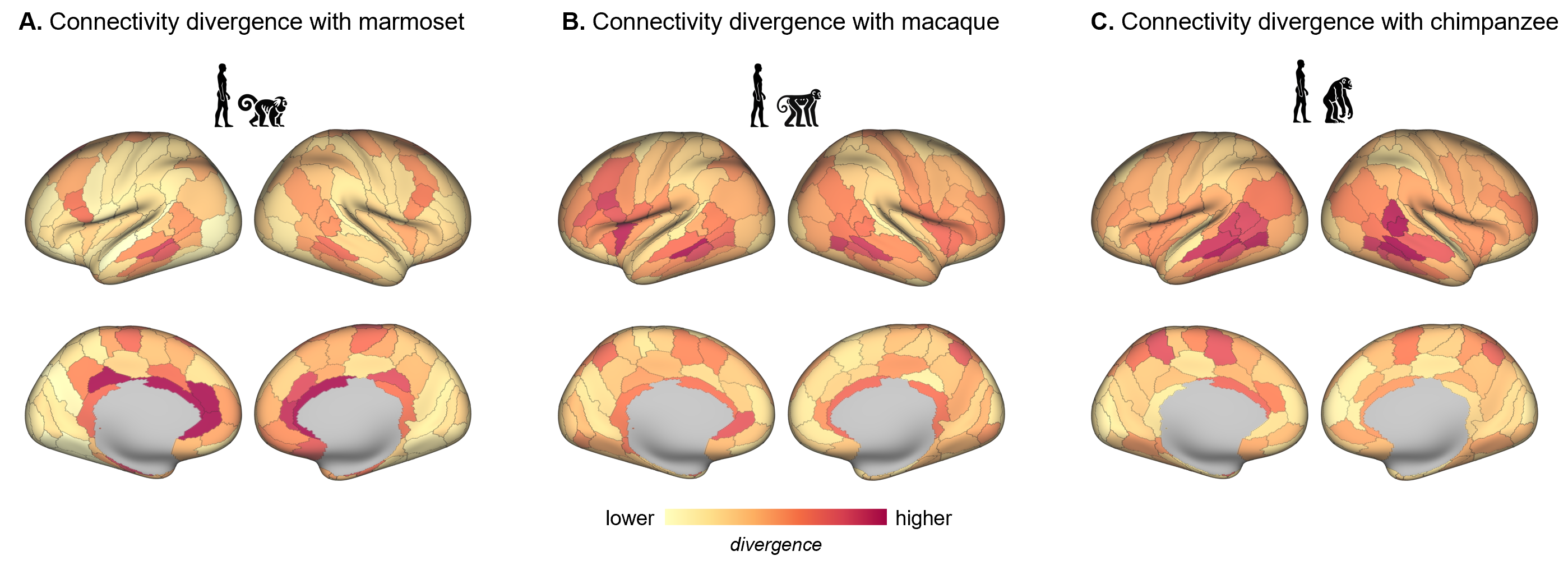

**Figure S4. Connectivity divergence between humans and (A) marmosets, (B) macaques, and (C) chimpanzees.**

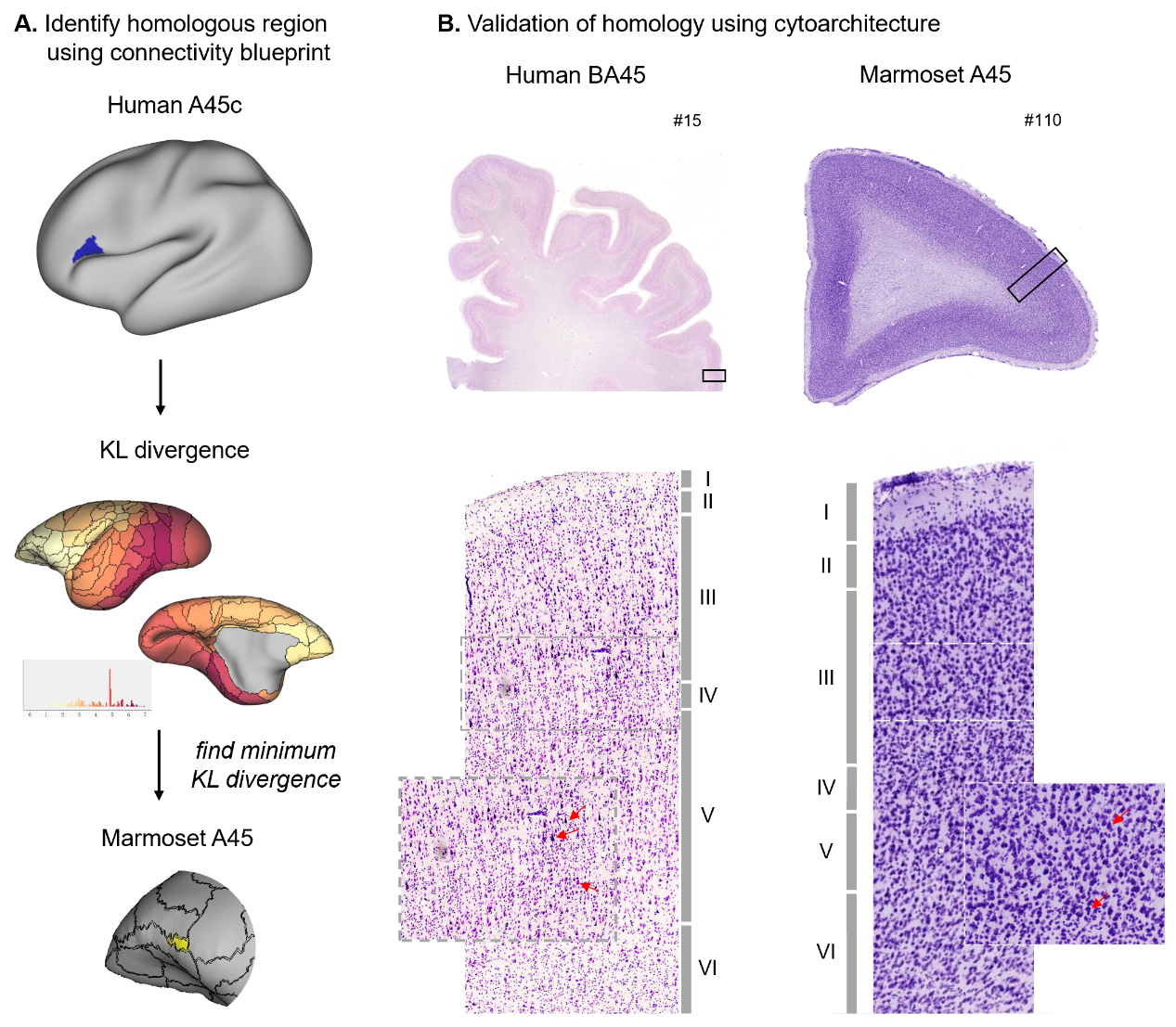

**Figure S5. Identification of homologous regions across species. (A)** Identification of homologous regions using connectivity blueprints. We used human A45c as an example. Calculating the KL divergence of the connectivity blueprint of human A45c and that of each marmoset subregions, we found the low divergence in the ventral frontal cortex. The minimum divergence was identified in area 45 in marmosets. **(B)** Validation of homology using cytoarchitecture. The cytoarchitectonic correspondence of area 45 in humans and marmosets was verified using Nissl-stained sections of this region from the two species. The similar features shared by the columns, including larger pyramidal cells (pointed by red arrows) in layer III and a well recognizable granular layer IV, indicated the cytoarchitectonic homology of area 45 between humans and marmosets.

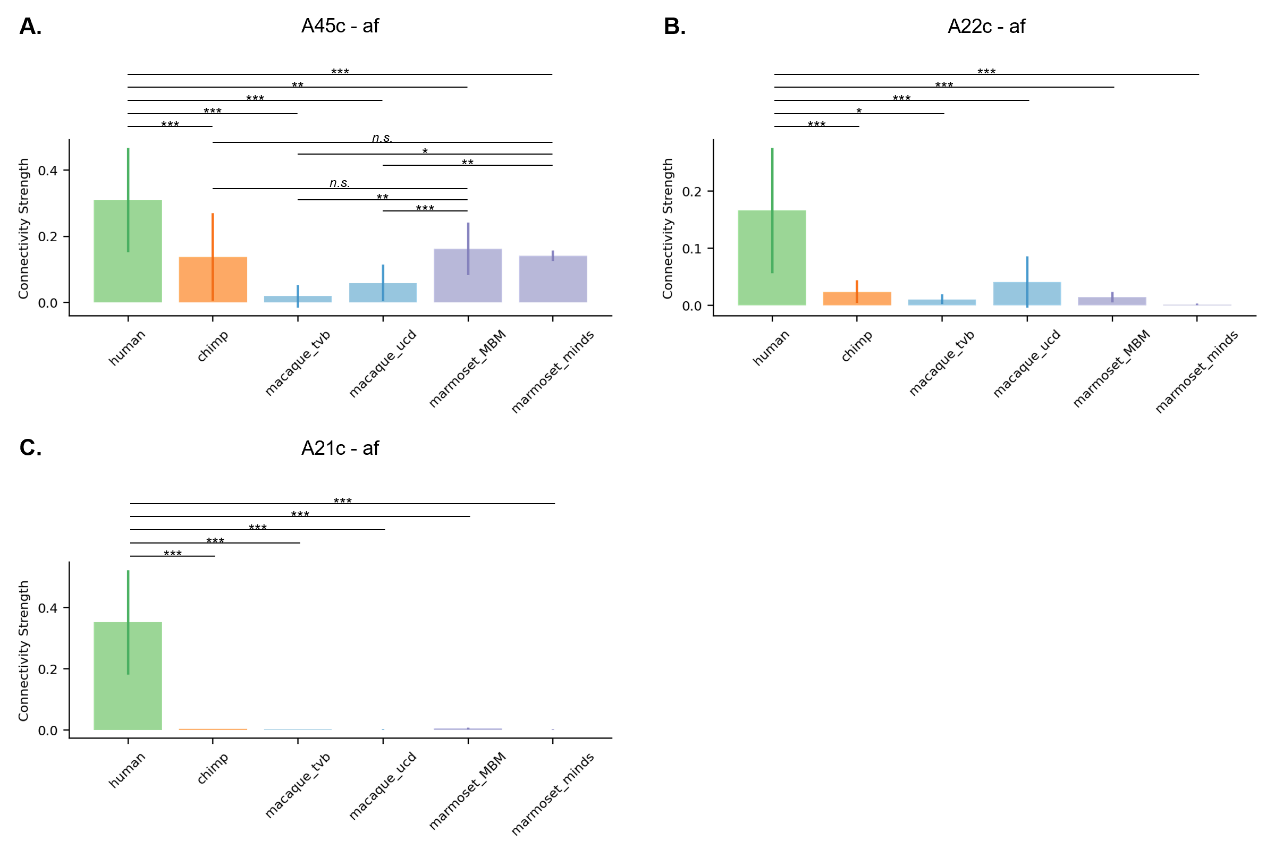

**Figure S6. Connectivity strength of af with subregions with high connectivity divergence. (A)** For caudal area 45 (A45c) in the ventral PFC, we found that its connection between the af was stronger in humans and marmosets than in macaques (*t*_human-chimp_ = 5.17, *p*_corrected_ < 0.001; *t*_human-macaque_tvb_ = 5.08, *p*_corrected_ < 0.001; *t*_human-macaque_ucd_ = 7.96, *p*_corrected_ < 0.001; *t*_human-marmoset_MBM_ = 4.19, *p*_corrected_ < 0.01; *t*_human-marmoset_minds_ = 8.10, *p*_corrected_ < 0.001; *t*_marmoset_MBM-chimp_ = 0.76, *p* = 0.45; *t*_marmoset_MBM-macaque_tvb_ = 4.85, *p*_corrected_ < 0.01; *t*_marmoset_MBM-macaque_ucd_ = 7.79, *p*_corrected_ < 0.001; *t*_marmoset_minds-chimp_ = 1.06, *p* = 0.29; *t*_marmoset_minds-macaque_tvb_ = 2.02, *p* < 0.05, uncorrected; *t*_marmoset_minds-macaque_ucd_ = 3.19, *p* < 0.01, uncorrected). **(B)** For caudal area 22c in the superior temporal gyrus, we found that its connection between the af was stronger in humans than other three species (*t*_human-chimp_ = 7.68, *p*_corrected_ < 0.001; *t*_human-macaque_tvb_ = 3.90, *p*_corrected_ < 0.05; *t*_human-macaque_ucd_ = 13.91, *p*_corrected_ < 0.001; *t*_human-marmoset_MBM_ = 6.62, *p*_corrected_ < 0.001; *t*_human-marmoset_minds_ = 8.10, *p*_corrected_ < 0.001). **(C)** For caudal area 21c in the middle temporal gyrus, we found that its connection between the af was stronger in humans than other three species (*t*_human-chimp_ = 13.52, *p*_corrected_ < 0.001; *t*_human-macaque_tvb_ = 5.70, *p*_corrected_ < 0.001; *t*_human-macaque_ucd_ = 8.83, *p*_corrected_ < 0.001; *t*_human-marmoset_MBM_ = 9.81, *p*_corrected_ < 0.001; *t*_human-marmoset_minds_ = 21.21, *p*_corrected_ < 0.001). *n.s.*, non-significant, * *p* < 0.05, ** *p* < 0.01, *** *p* < 0.001.

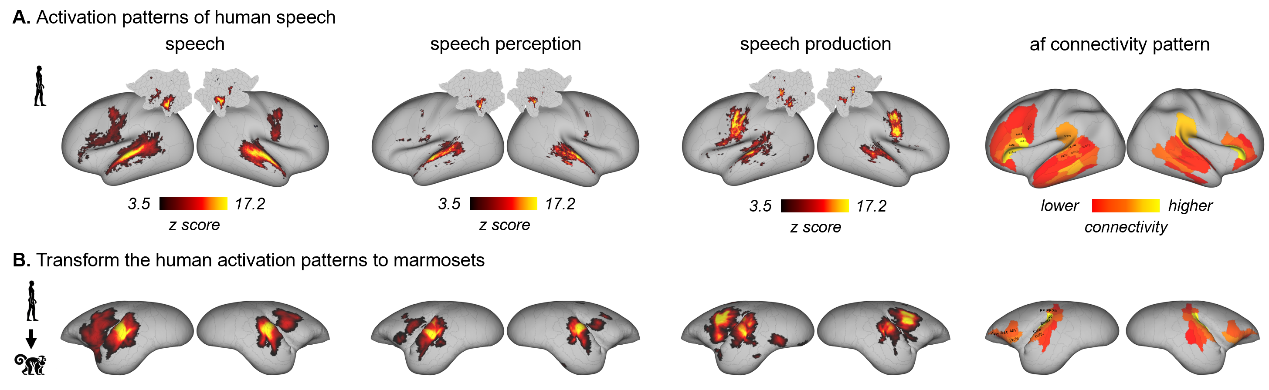

**Figure S7. Association between af and vocalization. (A)** Activation patterns of human speech and the connectivity patterns of af. **(B)** The transformed activation patterns in marmoset cortex.

### Supplementary tables

**Table S1. Scanning parameters of ultra-high-resolution diffusion datasets**

|  | Spatial resolution | Diffusion directions |
| --- | --- | --- |
| High-resolution marmoset data | 0.08 × 0.08 × 0.08 mm | 64 *b* = 2400 s/mm^2^  126 *b* = 4800 s/mm^2^ |
| High-resolution human data | 0.76 ×0.76 × 0.76 mm | 420 *b* = 1000 s/mm^2^  840 *b* = 2500 s/mm^2^ |
| High-resolution chimpanzee data | 0.5 ×0.5 × 0.5 mm | 55 *b* = 5000 s/mm^2^ |
| High-resolution macaque data | 0.2 × 0.2 × 0.2 mm | 30 *b* = 4000 s/mm^2^ |

**Table S2. Scanning parameters of lower-resolution diffusion datasets**

|  | Spatial resolution | Diffusion directions |
| --- | --- | --- |
| Marmoset dataset 1:  Marmoset Brain Mapping (MBM) | 0.5 × 0.5 × 0.5 mm | 64 *b* = 1000 s/mm^2^  128 *b* = 2000 s/mm^2^ |
| Marmoset dataset 2:  Brain/MINDS Marmoset Brain MRI Dataset (Brain/MINDS) | 0.35 × 0.35 × 0.7 mm | 30 *b* = 1000 s/mm^2^  60 *b* = 3000 s/mm^2^ |
| Human dataset 1:  Human Connectome Project (HCP) | 1.25 ×1.25 × 1.25 mm | 90 *b* = 1000 s/mm^2^  90 *b* = 2000 s/mm^2^  90 *b* = 3000 s/mm^2^ |
| Chimpanzee dataset 1:  National Chimpanzee Brain Resource (NCBR) | 1.8 ×1.8 × 1.8 mm | 60 *b* = 1000 s/mm^2^ |
| Macaque dataset 1:  TheVirtualBrain (tvb) | 1 ×1 × 1 mm | 64 *b* = 1000 s/mm^2^ |
| Macaque dataset 2:  UC-Davis macaque data (ucd) | 0.7 × 0.7 × 1.4 mm | 60 *b* = 1600 s/mm^2^ |
